# Benchmarking Pre-trained Genomic Language Models for RNA Sequence-Related Predictive Applications

**DOI:** 10.1101/2025.03.05.641574

**Authors:** Ningyuan You, Chang Liu, Hai Lin, Sai Wu, Gang Chen, Ning Shen

**Affiliations:** Department of Obstetrics and Gynecology of Sir Run Run Shaw Hospital & Liangzhu Laboratory, Zhejiang University School of Medicine, Hangzhou, Zhejiang, China; State Key Laboratory of Blockchain and Data Security Zhejiang University

## Abstract

RNA plays a pivotal role in diverse cellular functions across organisms. Developing computational algorithms for RNA sequence related questions is highly valuable. Recently, genomic language models (gLMs) with pre-training have emerged, offering flexibility for various downstream prediction tasks. However, comprehensive and fair evaluations of gLMs are lacking. In this study, we benchmark eight gLMs on prediction tasks covering four RNA processes, highlighting their strengths and limitations. While gLMs excel in performance overall, the larger model is not always better. Interestingly, models that integrate biological information consistently perform well in related tasks. Notably, gLMs demonstrate superior performance with limited training data, whereas task-specific methods achieve comparable performance with better computational efficiency when sufficient training data is available. Finally, we provide recommendations for model selection in different scenarios. These evaluation results underscore the potential of gLMs and suggest areas for future improvement.

## Introduction

RNA performs a remarkable range of molecular functions and involves cascades of biological processes to execute these functions^1^. The highly dynamic and orchestrated biological processes of RNA contribute to the phenotypic diversities of the cell. Compelling evidence has accumulated to demonstrate the causal role of dysregulated RNA processes in human diseases. For example, Amyotrophic Lateral Sclerosis (ALS) is a devastating disease caused by the degeneration of motor neurons. It has been reported to be associated with various RNA dysregulations, such as dysregulated microRNA biosynthesis^2^, globally reduced N6-methyladenosine (m6A) contributing to neurodegeneration^3^, and cryptic splicing due to TDP-43 depletion leading to the inclusion of erroneous exon^4^. As high-throughput experiments continue to expand our understanding of these RNA processes, there is a growing need for computational tools to analyze and interpret the massive experimental data collected regarding various aspects of RNA biology^5^.

Recent advancements in artificial intelligence (AI) have revolutionized RNA biology research. In particular, supervised deep learning models have gained prominence in tackling specific RNA-related tasks.^6–9^. For instance, DeepM6ASeq^10^ employs Convolutional Neural Network (CNN) architectures to predict sites of m6A modifications. Similarly, SpliceAI^11^ constructs a sophisticated CNN framework to identify splice sites within pre-mRNA sequences, analyzing flanking contexts up to 10 kilobases. More recent approaches, such as SpTransformer^12^, have been tailored to address distinct biological challenges by leveraging the Transformer architecture, which processes sequential data and dynamically assigns varying weights to different input segments. Each of these models is specifically designed for one or a set of particular tasks, contributing significantly to advancing biological research.

In parallel, pre-trained large language models (LLMs) based on Transformer architectures have achieved significant success across various Natural Language Processing (NLP) tasks, establishing a prominent trend in deep learning. Given the similarity between natural languages and biological sequences^13^, NLP techniques have been adapted to model biological sequence-related tasks, delivering impressive results in protein language modeling. For example, AlphaFold^14^ has made groundbreaking advances in predicting the 3D structures of proteins, revolutionizing structural biology. Similarly, ESM-2^15^ excels at generating meaningful protein sequence embeddings for diverse downstream applications. Inspired by the successes in protein modeling, genomic language models (gLMs) built on genomic sequences, including both DNA and RNA, have emerged to address tasks of RNA biology^16,17^.

As representatives of this trend, a new wave of pre-trained gLMs has been developed as powerful tools for studying RNA biology. Inspired by the Bidirectional Encoder Representations from Transformers framework in NLP^18^, these gLMs incorporate specific adaptations to address biological tasks. They serve as “foundation models” that can be flexibly adapted to a wide range of downstream applications^17^. These language models process sequences using “tokens” as fundamental units (e.g., words in natural language, codons for exons, or nucleotides for ncRNAs). In the training process, the models first undergo self-supervised training on large-scale, unlabeled genomic datasets. This step involves predicting the content of masked tokens within the biological sequences, enabling the models to learn the universal semantics of genomic sequence data. Beyond this self-supervised learning approach, a variety of model architectures, parameters, training datasets, and specialized techniques have been implemented to create diverse gLMs. For instance, RNAErnie^19^ is trained with biologically meaningful motifs. SpliceBERT^20^ integrates knowledge from multiple vertebrates. DNABERT2^21^ introduces irregular tokenization based on frequently occurred nucleotide pairs in the genome. More recently, Nucleotide Transformer^22^ has introduced a large-scale architecture designed to handle extended sequence contexts, enhancing its performance in complex prediction tasks. Leveraging the power of deep learning alongside large and diverse biological perspectives, these models have offer profound insights into challenging bioinformatics problems in RNA biology and research.

While the development of various deep-learning models has significantly advanced our understanding and modeling of diverse RNA processes, fair and comprehensive comparisons of these models are lacking. Persistent questions include whether gLMs as foundational models can effectively address all RNA-related tasks, whether they consistently outperform deep-learning algorithms tailored to specific RNA processes, and whether a definitive leader among foundational gLM models exists for all RNA-related tasks. To address these questions, we developed a benchmark framework to systematically evaluate the performance of gLMs across multiple RNA biological process tasks. Surprisingly, while results highlight the substantial potential of gLMs in advancing RNA biology, for model performance, the larger is not always better. Variations in training input and tokenization strategy may confer performance advantage for different gLMs in specific RNA-realted tasks. Addtionally, our benchmark show that lightweight, specialized methods achieve comparable performance at much lower computational cost when reasonably sufficient training data is available. Finally, we provide more specific discussion and guidance about the recommendations of models to use, taking into account the trade-off between performance versatility and computational efficiency. Our findings provide insights into the future development of more powerful gLMs, and offer practical guidance for researchers in selecting the most suitable deep-learning models or training strategies for their specific RNA biology question.

## Results

### Performance measurement of gLMs across various RNA sequence-related tasks

In this work, we implemented a flexible and extensible framework to evaluate multiple pre-trained gLMs on tasks related to RNA sequences (Fig. 1). To comprehensively benchmark these models, we concentrated on four representative sequence-related tasks in RNA biology: ncRNA function classification, m6A modification prediction, alteration splicing prediction, and translation efficiency prediction. These tasks are fundamental in RNA biology and have been subjects of intense research, with the development and application of both task-specific and gLM-based models in recent years. Moreover, these tasks illustrate diverse computational problems in RNA sequence-related analyses. Specifically, ncRNA classification is a sequence-level multi-classification task that requires models to recognize the overall characteristics of the whole input sequences. The m6A modification classification is a binary classification task, where models classify the central position of the input sequence based on its sequence context. The splice site prediction is a nucleotide-resolution task, where models are required to provide predictions for each position of a sequence, rather than treating the sequence as a whole. Lastly, the translation efficiency prediction represents a regression task that requires the models to predict the mean ribosome loading (MRL) for given RNA sequences. For each RNA process task, we selected a widely accepted representative dataset for benchmarking, which encompasses training sample sizes from thousands to hundreds of millions (Fig. 1a). The task design and benchmark datasets ensures a robust and comprehensive evaluation of model performance across various situations.

**Figure 1.**
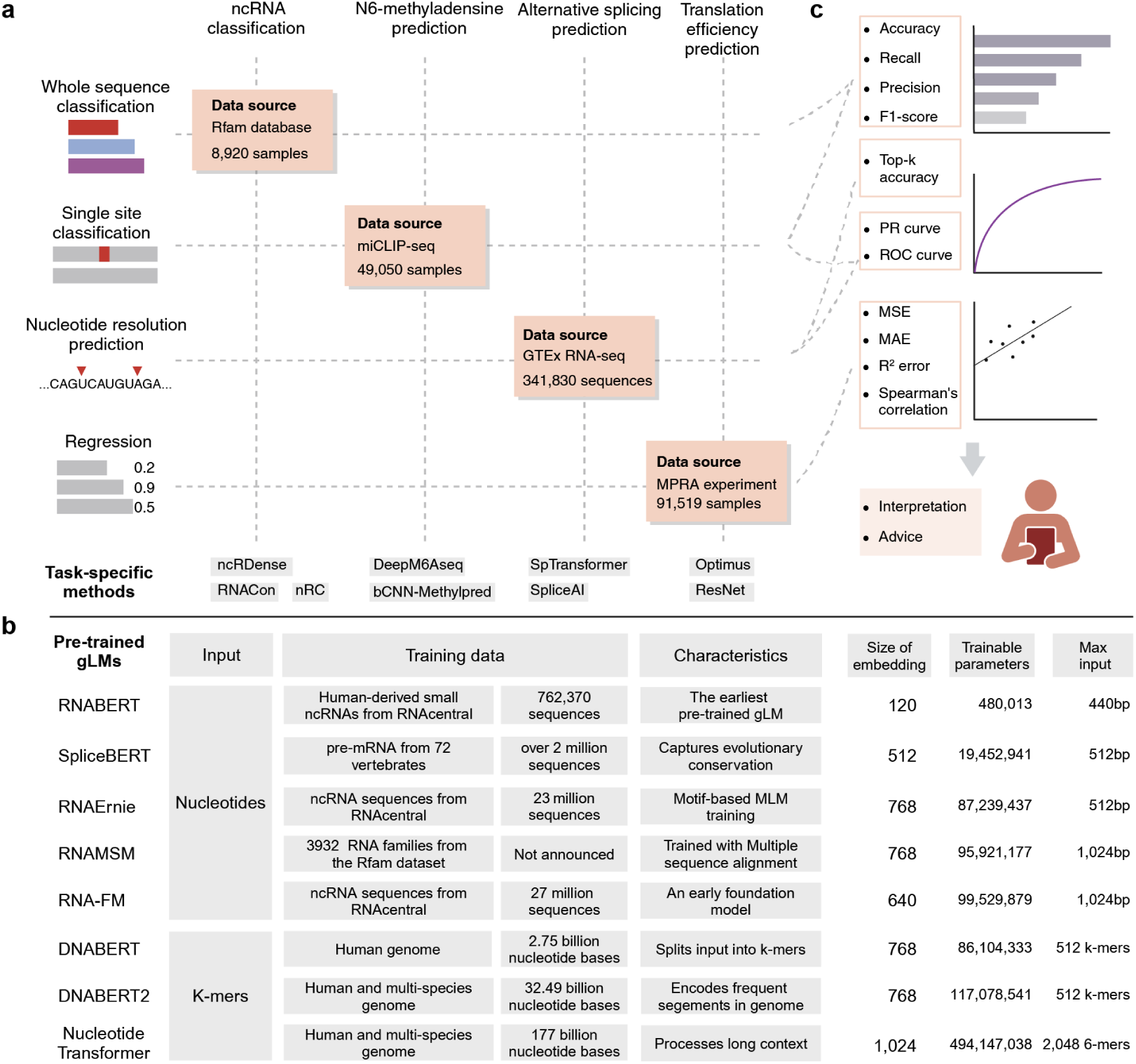
Schematic overview of benchmarking pre-trained gLMs in multiple RNA sequence-related tasks. **a** Overview of four typical biological tasks, showing the datasets and task-specific methods used in each task. **b** Model characteristics, specifications and training strategy of pre-trained gLMs. **c** Multiple metrics used to evaluate the models in each task.

We included eigth pre-trained gLMs with diverse model designs and training strategies for comparison. (Fig. 1b). These models differ in their training inputs, embedding sizes, number of parameters, and tokenization approaches. Specifically, RNABERT^23^ is the earliest attempt at a pre-trained BERT model that learns sequence features from human-derived small ncRNAs. It has significantly fewer trainable parameters than others. SpliceBERT^20^ employed primary RNA sequences of multiple vertebrates during pre-training. RNA-FM^24^ was developed as a foundational model designed to handle all non-coding RNA (ncRNA) sequences. It was an early attempt to employ massive unannotated sequences and a large model for pre-training. RNAErnie^19^ was built upon the Enhanced Representation through Knowledge Integration (ERNIE) framework. This framework is trained with unannotated sequences, and incorporates a specialized masking strategy that emphasize on continuous subsequences and well-known motifs, thereby enhancing the learning of RNA sequence features. RNAMSM^25^, inspired by the MSA-Transformer^26^, employed a specialized sequence-alignment strategy to train the model with RNA homologs from the Rfam dataset. DNABERT^27^ was trained with DNA data. It encodes input sequences by k-mers of fixed lengths, thereby incorporating motif-based sequence features. DNABERT2^21^ builds upon this foundation by refining the segmentation method.

Instead of using fixed-length k-mers, DNABERT2 splits the input sequences into irregular segments based on frequent pairs of nucleotides and genomic segments. In addition, it also uses multispecies genomes in training. Nucleotide Transformer^22^ surpassed other models both in training datasets and model scales. It uses a massive transformer model in order to consider the complex long-range context in the input sequence, thus offering enhanced performance in complex prediction tasks.

We adopted the officially released weights of the pre-trained gLMs and fine-tuned them for each specific task. For task-specific methods, we retrained the models using the selected benchmark datasets, except in cases where specialized training data was an integral part of the model design (see Methods). In addition, we incorporated several task-specific methods as benchmarks. These methods were developed to address prediction tasks for individual RNA processes. As representative task-specific models, we included ncRDense^28^, nRC^29^ and RNACon^30^ for non-coding RNA (ncRNA) classification, DeepM6ASeq^10^ and bCNN-Methylpred^31^ for N6-methyladenosine (m6A) modification prediction, SpliceAI^11^ and SpliceTransformer^12^ for splice site prediction, and ResNet^32^ and Optimus^33^ for MRL prediction, respectively. All evaluated models underwent a consistent data split strategy to avoid data leakage (Method).

To ensure a fair and comprehensive evaluation, we employed standardized metrics tailored to the specific characteristics of each task (Fig. 1c). For classification tasks, we differentiated between multi-class and binary scenarios. In the ncRNA classification task, as a multi-class problem, we focused on the model’s ability to distinguish among multiple categories. Thus, we used accuracy, precision, recall, and F1-score to evaluate overall performance. For the m6a prediction task, a binary classification task with balanced dataset, we employed PR-AUC and ROC-AUC to assess classifier performance across various thresholds. For the splicing prediction task, which requires nucleotide-level resolution, the extreme class imbalance posed a significant challenge due to the sparse distribution of splice sites within pre-mRNA sequences. Traditional accuracy metrics are insufficient for such tasks. Thus, we employed top-k accuracy and PR-AUC to measure the model’s ability to identify accurate splice sites. Finally, for the MRL prediction task, where quantitative predictions are essential, we used MSE, MAE, R-squared error, and Spearman’s correlation to evaluate the prediction performance. The detailed computation of these metrics is described in the Methods section.

In addition to benchmark the performance of these models, we also delved into the characteristics that confer their practical utility in addressing these tasks. To better understand the performance strength of various models, we have conducted a series of analyses, including data down-sampling and ablation studies, across varying levels of difficulty. Additionally, we have profiled the computational time cost associated with each model, thereby providing a comprehensive evaluation of both efficiency and effectiveness.

### Classification of non-coding RNAs

The ncRNAs encompass diverse classes and are reported to account for 76-97% of the human genome^34^. These ncRNAs are short nucleic acid sequences that do not translate into proteins. Different categories of ncRNAs vary in terms of sequence length, folding patterns, and biological functions^35^. Previous studies have demonstrated that ncRNAs are involved in key molecular functions and contribute to specific biological processes and phenotypes^36,37^. Despite the substantial number of ncRNAs identified to date, the understanding of their characteristics and functions remains limited^35,38^. Accurate computational classification of ncRNAs may provide biologists with valuable insights into the roles of ncRNAs in gene regulation, cellular processes, and disease mechanisms. Consequently, the proper identification and classification of ncRNAs have emerged as a significant challenge in the field of bioinformatics.

To evaluate the ability of gLMs to recognize RNA sequence features, we set up a multi-class classification task that aimed at distinguishing non-coding RNA (ncRNA) sequences. The labeled dataset for this task, named nRC, was constructed based on the Rfam database release 12^39^. This dataset consists of a balanced collection of sequences, comprising 6,320 training sequences and 2,600 testing sequences, each labeled with one of 13 classes. The sequence lengths range from 39bp to 512bp. We fine-tuned every gLM on this dataset using RNA sequences as input. Additionally, we included three task-specific methods, i.e., nRC, ncRDense, and RNACon, for benchmarking. Unlike gLMs, which rely solely on sequence data, these methods incorporate additional features as input. RNACon^30^ manually extracts structure-based sequence features using multiple bioinformatics tools and employs a RandomForest classifier to classify ncRNAs. nRC^29^ calculates graph-based structural features and uses a small CNN for final classification. Concordantly, ncRDense^28^ implements a deep CNN architecture with multiple CNN blocks to process predefined features such as sequence, secondary structure, and electron-ion interaction pseudopotentials. Further details on data processing and model fine-tuning are provided in the Methods section.

The ncRNA classification task requires the models to analyze the features of the entire input sequence and classify it into one of multiple classes (Fig. 2a). The model performance was evaluated across several metrics, including accuracy (Fig. 2b), precision (Fig. 2c), recall (Fig. 2d), and F1-score (Fig. 2e). Overall, the gLMs demonstrated superior performance compared to earlier task-specific methods, with the exception of the earliest gLM, RNABERT. Among all gLMs, RNA-FM achieved the highest performance in all evaluation metrics. Notably, SpliceBERT also achieved high performance but with a significantly reduced computational cost (Fig. 2b, Supplementary Fig. 1). Although initially designed for splicing-related tasks, this outcome was expected, as SpliceBERT learned general RNA-related knowledge during its pre-training stage with pre-mRNA data. DNABERT and DNABERT2, both pre-trained on DNA datasets, achieved moderate performance on this task, suggesting their ability to extract RNA-related features during pre-training. Intriguingly, despite DNABERT2 featuring a larger training dataset and increased model complexity compared to its predecessor, it demonstrated lower performance. A distinctive feature of DNABERT2 is its unique approach to tokenization approach, which segments sequences based on frequency across the genome. While this tokenization strategy helps the model to capture sequence motifs, it is possible that the frequently occurring sequence motifs may not be the most informative to distinguish the ncRNA classes. Instead, this approach might cause the model to focus on less relevant motifs, potentially impeding its overall performance. Concordantly, Nucleotide Transformer, developed with huge number of training samples and parameters, exhibited moderate performance in this task. This suggests that larger models did not confer learning advantage in this task.

**Figure 2.**
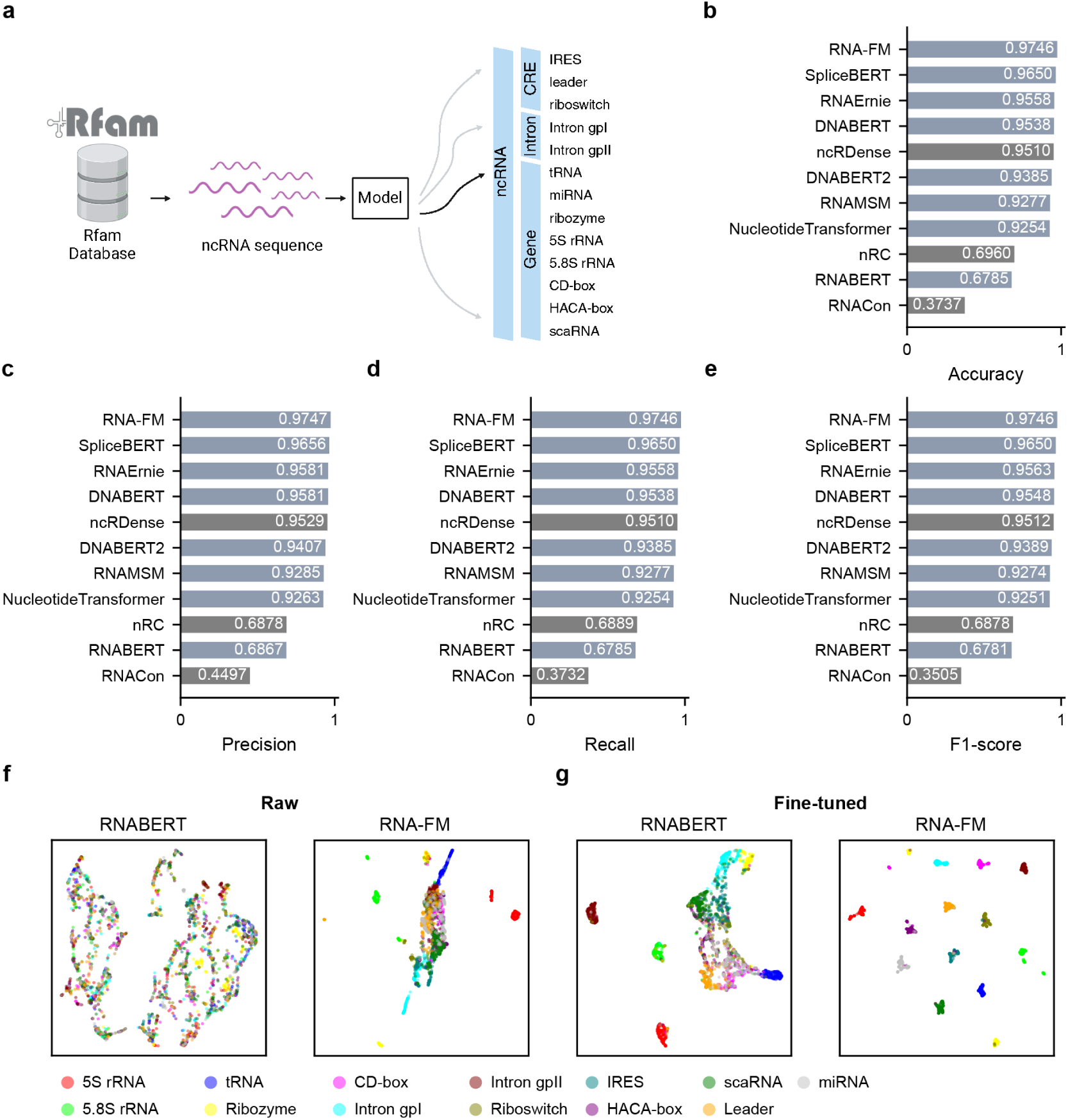
Sequence-level multi-classification task of ncRNA classification. **a** Diagram of this task: Classifying an input sequence into 1 of 13 ncRNA predefined categories. **b-e** Multiple measurements comparing across different models: **b** accuracy, **c** precision, **d** recall, and **e** F1-score metrics. The task-specific methods were marked with gray color. **f-g** UMAP visualization of sequence embeddings extracted by two representative models, RNABERT and RNA-FM, before (**f**) and after (**g**) fine-tuning.

Importantly, the task-specific model ncRDense^28^ also achieved impressive performance across all evaluation metrics, with only a 0.02 difference in performance from the best model. Key strength of ncRDense include its ability to incorporate RNA structural information provided by other bioinformatics tools as additional input and its use of an advanced dense CNN architecture. In contrast, despite incorporating structural features, the RNACon method, based on Random Forest, and the nRC method, which employed a simpler CNN model, exhibited significantly lower performance. This result underscores the superior capability of complex deep-learning architectures in predicting the ncRNA classes.

Despite the generally comparable performance of most pre-trained gLMs, RNABERT demonstrated limited capability on this task. The primary difference between RNABERT and other gLMs lie in the scale of their training datasets and the number of trainable parameters (Fig. 1a). As one of the earlier pre-trained models, RNABERT has a significantly simpler deep-learning architecture, which may limit its effectiveness across tasks. We further conducted a visualization analysis of the sequence embeddings extracted by the two models: RNA-FM and RNABERT. The models showed varying abilities to aggregate ncRNA sequences through unsupervised methods (Fig. 2f). RNA-FM demonstrated a distinct advantage both before and after fine-tuning for this task, highlighting the importance of evaluating the unique characteristics of different pre-trained gLMs for specific biological research applications.

The efficiency of learning and inference was also evaluated (Supplementary Fig. 1a). Pre-trained gLMs exhibited varying rates of task acquisition. For example, SpliceBERT could be trained three times faster than RNA-FM. Despite its smaller size, SpliceBERT proved sufficient for handling most aspects of this classification task. In contrast, larger models required substantially more computational resources for a minor improvement in performance. Moreover, when subjected to the same number of training epochs, the models exhibited different learning dynamics (Supplementary Fig. 1b). RNA-FM and RNABERT displayed stark contrasts in learning speed under identical training settings. RNA-FM quickly grasped the task and achieved optimal performance within fewer than 10 epochs. In contrast, RNABERT initially struggled to understand the task and learned at a slower pace compared to the other models. These findings are consistent with the visualization analysis in Figures 2f-g.

In summary, while there were notable differences in computational efficiency and learning speed, the majority of the gLMs performed well. Intriguingly, for gLMs, the larger models are not better. On the other hand, among the task-specific models, only ncRDense, which was built on a dense CNN architecture and incorporates additional biological features, demonstrated superior performance, underscoring the value of complex deep-learning architecture in such prediction tasks.

### Prediction of N6-methyladenosine related with sequence context

N6-methyladenosine (m6A) is the most prevalent and evolutionarily conserved modification of polymerase II transcribed RNAs^40^. Found in nearly all types of RNAs, m6A is present across a wide range of organisms, from bacteria to animals^40,41^. This modification plays a crucial role in various RNA activities, including alternative splicing, microRNA interactions, and RNA translation efficiency^10^. High-throughput experimental technologies, such as Methylated RNA Immunoprecipitation Sequencing (MeRIP-Seq)^42^ and m6A Individual-nucleotide-resolution Cross-linking and Immunoprecipitation Sequencing (miCLIP-Seq)^43^, have been instrumental in identifying m6A modification sites. Based on these experimental data, several computational methods have been developed, achieving high accuracy in predicting m6A sites^10,31^.

Predicting m6A modification can be framed as a binary classification task, where the goal is to evaluate whether the central nucleotide in a sequence undergoes m6A modifications. We selected the human miCLIP-Seq dataset, compiled and pre-processed by Linder et al.^43^, for benchmark evaluation. This dataset consists of 49,050 training samples and 12,611 testing samples, with each input sequence containing 101 nucleotides. The models were trained to predict whether the central nucleotide of each input sequence would undergo m6A modification (Fig. 3a). For comparison, two deep-learning methods were also included in the analysis. DeepM6ASeq^10^ combines CNN layers with an LSTM network to process one-hot encoded RNA sequences, while bCNN-Methylpred^31^ employed multiple CNN blocks as “branches” to process input sequences with manually derived encoding strategies. The encoding for bCNN-Methylpred includes one-hot encoding, frequent nucleotide pairs, and chemical properties. The time cost and training curves for all methods are provided in Supplementary Fig. 2.

**Figure 3.**
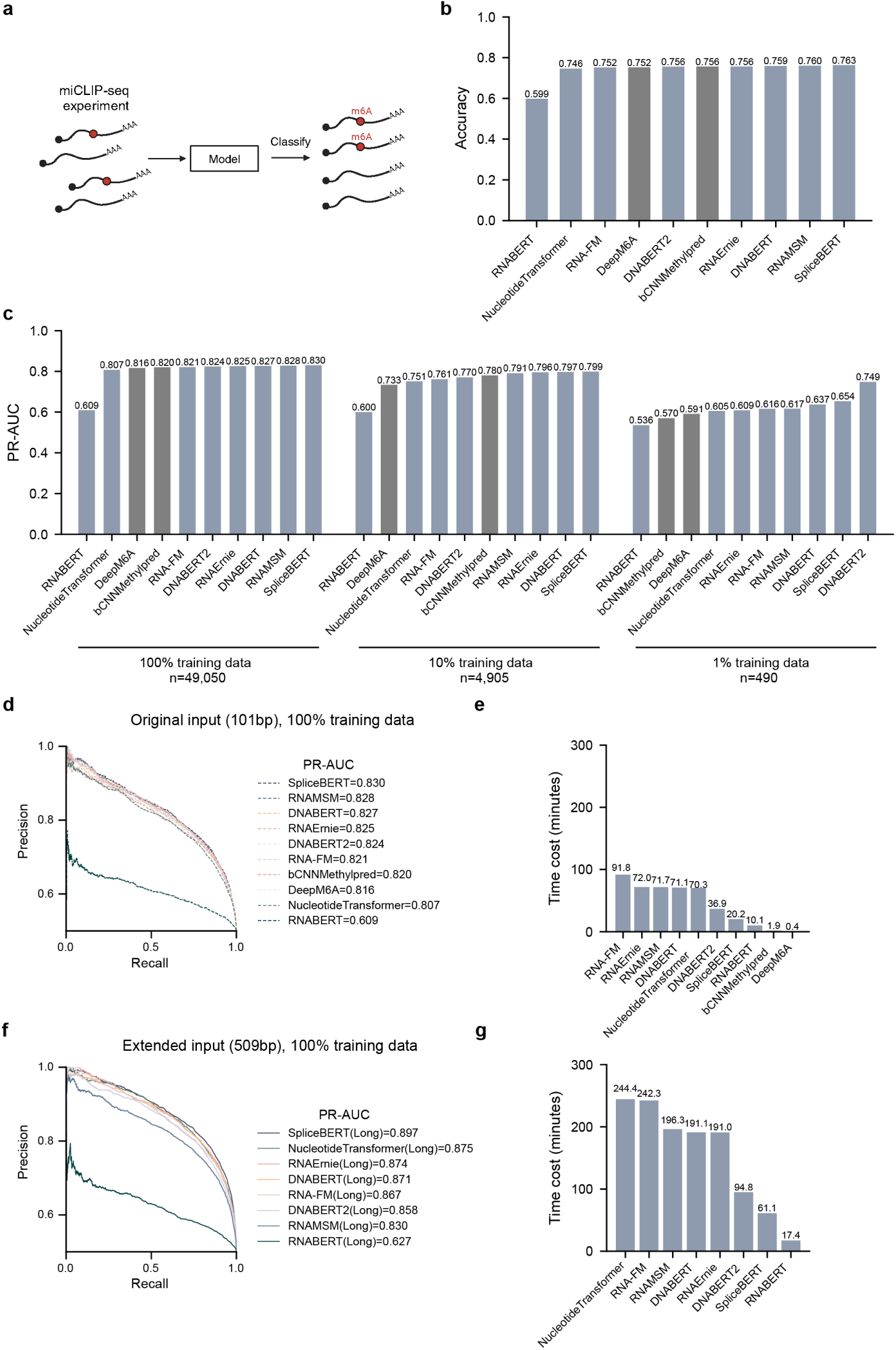
Classification task of predicting m6A modification. **a** Diagram of this task: Distinguishing whether the center of each input sequence undergoes m6A modification. **b** Evaluation of the prediction accuracy across different models. **c** Performance of models when different scales of training data were provided. **d,e** PR-Curve and fine-tuning time for models on the original dataset. **f,g** PR-Curve and fine-tuning time for models when extended lengths of input sequence were given.

Benchmark results show that most of the gLMs slightly outperformed the task-specific methods in predicting m6A modifications (Fig. 3b). Among the gLMs, SpliceBERT achieved the highest performance in all evaluation metrics. As previously described, SpliceBERT was pre-trained on primary RNA sequences from multiple vertebrates. Given that m6A modifications exhibit a certain degree of evolutionary conservation across species, this pre-training strategy likely contributes to its superior performance on this task. In contrast, RNABERT, as an earlier model with a significantly smaller parameter scale, was the only model that performed notably worse than all the other models.

Importantly, the performance scores among gLMs and task-specific models were quite similar. While the accuracy and PRAUC of the best-performing model, SpliceBERT, were 0.763 and 0.830, respectively, the performance scores for the task-specific model DeepM6ASeq were slightly lower, at 0.752 and 0.816. We then compared the training efficiency of all the models and found that DeepM6Aseq and bCNN-Methylpred can achieve training and application speeds up to 50 times faster than the leading gLM method (Supplementary Fig. 2a). Thus, the above task-specific methods are generally recommended over gLMs considering the balance between performance and efficiency.

Next, we investigated the effect of sample size on model performance. Results showed that pre-trained gLMs demonstrate more superior performance advantage compared to the task-specific models under a limited sample size (Fig. 3c). Specifically, when 100% of the training data (49,050 samples) were used, all methods achieved similar performance. In this scenario, the gap between bCNN-Methylpred and the best-performing gLMs was approximately 1%. However, with 10% of the original training data used, a more substantial difference emerged among the models. When the training data were further reduced to only 1%, the best-performing gLMs achieved a performance advantage of about 16% over the task-specific models, highlighting the superior performance of pre-trained models in situations with a small sample size. Interestingly, the motif-based gLM, DNABERT2, demonstrated a clear performance advantage over all other methods when only 1% of the original training data was used. As mentioned above, DNABERT2 employs a specialized tokenization strategy to partition input sequences into segments. This approach may enable the model to efficiently capture cis-regulatory elements, especially in scenarios where training data is limited.

Furthermore, considering that pre-trained gLMs are designed to extract contextual information over long range sequences, we conducted a specialized ablation study. Specifically, we increased the length of each input sequence from 101 nucleotides to 509 nucleotides by incorporating flanking sequences from the hg38 reference genome (note that most models support a maximum input length of 510-512 nucleotides; see Methods section). This adjustment provided additional contextual information for the larger gLM models. The extended models are denoted with a “(Long)” suffix, e.g., RNAFM(Long), RNABERT(Long), etc. As expected, all gLMs with longer inputs demonstrated superior performance compared to their original versions (Fig. 3d, Supplementary Fig. 3). Simultaneously, the performance disparity among gLMs was strengthened. Nucleotide Transformer demonstrated substantial improvements when processing extended input sequences (Fig. 3f, Supplementary Fig. 3). Its PR-AUC increased by 7%, securing the second position in the performance rankings. Similarly, RNAErnie and RNA-FM exhibited marked enhancements with longer input sequences. SpliceBERT remained the winner among all gLMs. Notably, processing longer input sequences required substantially more computational time for all models.

Together, the majority of the gLMs performed well, with similar performance scores compared to task-specific deep-learning models. Pre-training strategy on genome data from multiple vertebrates provided SpliceBERT performance advantage in this task, likely due to information gained from evolution conservation. Additionally, pre-trained gLMs demonstrated a clear advantage over task-specific models when limited training data was provided. They can also benefit from longer sequence contexts to achieve better performance.

### Prediction of alternative splicing sites across tissue types

Splice sites, nucleotide sequences that define the junctions between exons and introns in pre-mRNA, are essential for the precise removal of introns and the ligation of exons during the splicing process^44^. Splice sites are crucial in alternative splicing, determining the variability of mRNA transcripts produced from a single gene. More than 90% of human genes undergo this essential biological process.^45^. Disease-causing mutations in the splice sites or cis-regulatory elements of splicing are recognized as the second most common mechanism in genetic diseases^46^. Therefore, accurate prediction on splice sites is critically important for both biological research and medical genetics.

Mutations in pre-mRNA sequences can affect splicing over a broad region surrounding the original splice sites^47^. For example, *TP53* c.375G>T mutation have been reported to activate a cryptic splice site 200bp upstream of the original splice site, leading to an in-frame deletion of codons^48^. Thus, computational methods predicting all potential splice sites within an extended input sequence is helpful to directly interpret splicing effects of mutations. However, previous gLM studies, such as DNABERT2 and SpliceBERT, treated splice site prediction as a sequence-level classification task, where models output a single score for the entire input sequence to identify whether the center of the sequence is a splice site^20,21^. This approach limits their practical applications for downstream analysis. To address this issue, we introduced a nucleotide-resolution classification task that necessitates models to predict all splice sites in an interval of the pre-mRNA sequence. Moreover, alternative splicing is highly dynamic and varies across cell types and tissues^49^, which has led to increasing attention on tissue-specific splicing prediction in recent years^50–52^. Therefore, we compiled three tasks with varying degrees of challenge to benchmark splicing prediction, using the human adult tissue splicing dataset from the GTEx database^53^ (Fig. 4a; Methods).

**Figure 4.**
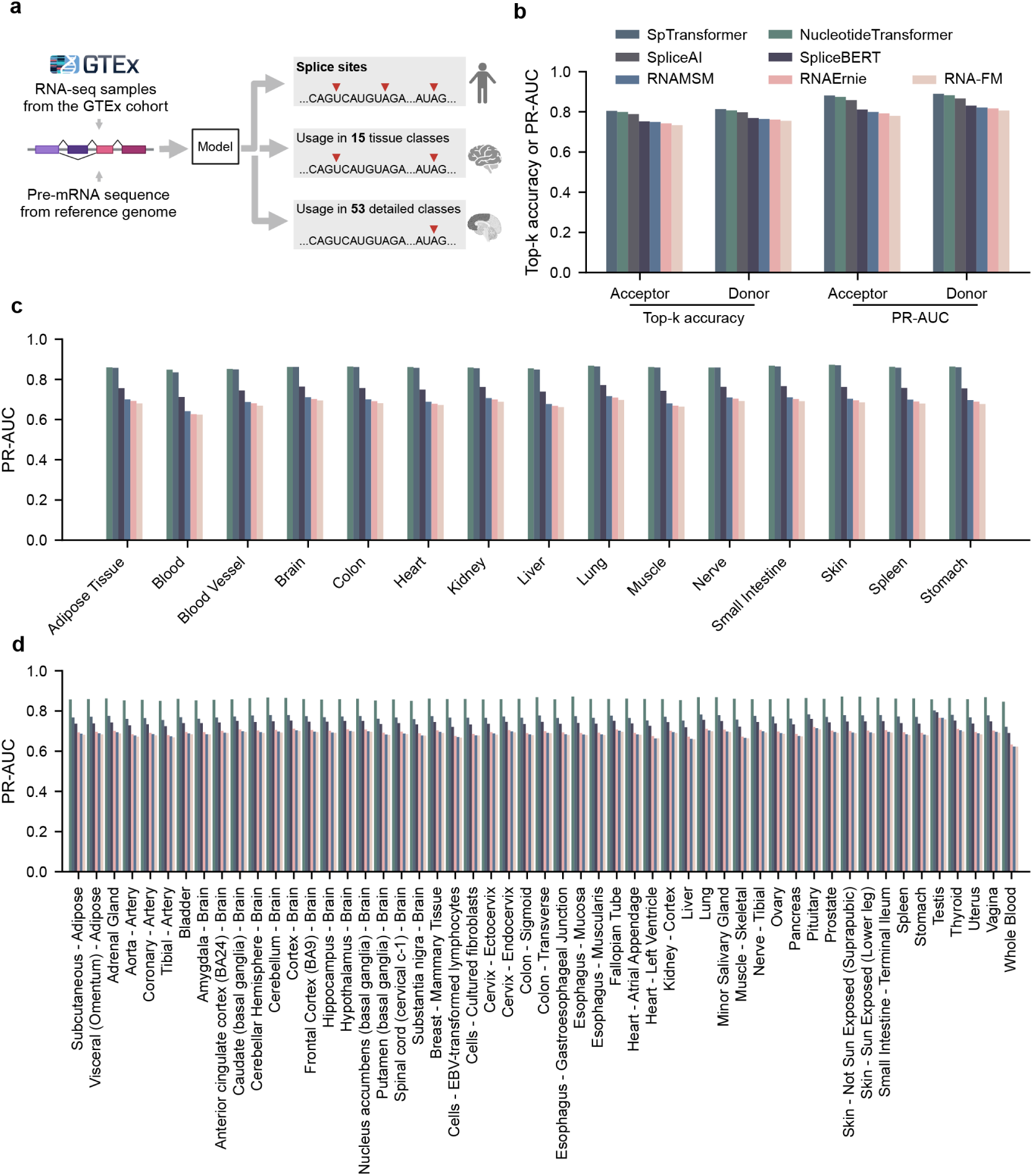
Nucleotide-level task of alternative splicing prediction. **a** Splicing datasets generated on three different levels. The 15 human tissue categories were defined using the “SMTS” variable (“Tissue Type, area from which the tissue sample was taken”), while the 53 tissue subtypes were defined using the “SMTSD” variable (“Tissue Type, more specific detail of tissue type”), according to annotations from the GTEx cohort. **b** Performance of models on the site type classification task. **c** Prediction of splice sites usage across 15 human tissues by different models. **d** Prediction of splice sites usage across 53 tissue subtypes by different models.

Previous task-specific methods, such as SpTransformer and SpliceAI, were designed to score all nucleotides within a range of 1,000 to 5,000 nucleotides simultaneously^11,12^. In contrast, pre-trained gLMs typically support input sequences ranging from 440 to 1,024 nucleotides. To balance model requirements with practical applications, we divided the input data into 500bp sequence for all methods. We included a total of 242,017 training sequences, comprising 121,008,500 nucleotides, and 32,773 testing sequences, comprising 16,386,500 nucleotides.

In splicing benchmark, the first task was to classify each position of the 500bp input sequence as one of 3 classes: splice acceptor, splice donor, or neither. The second challenge involved distinguishing different usages of splice sites across 15 human tissues, adding prediction output with 15 dimensions for splicing evaluation. Finally, we benchmarked the models using 53 tissue subtypes as labeled by GTEx, further challenging the models with increasing output dimensionality.

In these benchmark tasks, several gLMs, such as DNABERT, DNABERT2, and RNABERT, were incompatible with this nucleotide-level task and thus were excluded from this comparison (Methods). Two deep models specialized for splicing prediction, SpliceAI^11^ and SpTransformer^12^ were included for comparison. SpliceAI is a CNN-based deep model, while SpTransformer is a Transformer model that does not undergo self-supervised pre-training like gLMs. Both models were specifically designed to incorporate context features of up to 10,000bp. In contrast, the gLMs can only process input sequences of 512 or 1,024bp, due to limitations in model structure and current GPU technology.

The task-specific method SpTransformer and the pre-trained gLM Nucleotide Transformer achieved the highest performance in splice site prediction (Fig.4b) and the tissue-specific splicing prediction tasks (Fig. 4c). SpliceAI achieved third performance in splice site prediction tasks but was excluded from the other two tasks due to its missing function for tissue usage prediction. In the first task, SpliceBERT and other gLMs maintained comparative top-k accuracy and PR-AUC to the top models (Fig.4b). However, in the more challenging prediction tasks for 15 human tissues, the performance gap between methods became more pronounced (Fig.4c). Surprisingly, in the most complex task for 53 subtypes, Nucleotide Transformer exhibited a clear advantage in PR-AUC, while the performances of all the models dropped as the task complexity increased. Notably, Nucleotide Transformer maintained consistent PR-AUC values at about 0.85 among all 3 tasks. In contrast, SpTransformer struggle with the most complex task, achieving PR-AUC values below 0.77 across 53 tissue subtypes.

When evaluating the computational efficiency, SpliceAI, as a CNN model, demonstrated significantly faster processing time compared to the gLMs (Supplementary Fig. 4a). SpliceBERT exhibited a distinct advantage over all other gLM-based approaches in terms of time performance. Although Nucleotide Transformer and SpTransformer achieved high scores on the evaluation metrics, they were the slowest methods among all evaluated models. There are two key distinctions between the well-performing and poorly-performing methods: 1) SpTransformer and SpliceAI were trained exclusively on splicing datasets and did not require the inclusion of other RNA-sequence-related features. 2) Nucleotide Transformer, SpTransformer, and SpliceAI could process input sequences up to 9,000bp, which is significantly longer than 512bp processed by other models. This result aligns with the well-established fact that cis-acting regulators of splicing events can be located remotely from the splicing sites. Unfortunately, these compared gLMs require computational resources on the order of *O*(*n*^2^) for both time and memory complexity when the input sequence contains n tokens. Extending the input sequence length from 512 bp to 9,000 bp results in an approximately 300-fold increase in GPU resource costs.

Given that these models were pre-trained using 4 to 8 high-end GPUs, processing longer sequences would necessitate either hundreds of GPUs or a corresponding increase in computational time, which is prohibitively expensive for most laboratories. Consequently, due to limitations in both model architecture and current GPU technology, these gLMs are currently unable to efficiently handle ultra-long input sequences. Nucleotide Transformer mitigates this issue by introducing a non-overlapping 6-mer encoding scheme, reducing the number of required tokens by a factor of sixth (i.e., 1,500 input tokens for 9,000 bp). This approach is followed by specialized prediction layers that decode prediction scores for each nucleotide. Additionally, Nucleotide Transformer employs Low-Rank Adaptation (LoRA), a fine-tuning strategy enabling efficient fine-tuning with only one GPU and a few days of processing. However, even with these optimizations, the process still require significantly more time compared to task-specific methods, and the initial pre-training remains computationally intensive, demanding substantial time and GPU resources.

Additionally, we conducted an ablation study to investigate the impact of varying training data sizes (Methods). In a reduced dataset using only approximately 25% of the training data, model performance declined to varying degrees. Specifically, the top-k accuracy of SpTransformer decreased by approximately 8%, while SpliceAI’s performance dropped by only 2%. Notably, the performance of pre-trained gLMs declined by about 1% to 3%, positioning Nucleotide Transformer and SpliceAI as the top two methods in this evaluation (Supplementary Fig.5). These findings are consistent with our previous understanding that Transformer-based models generally require more training resources compared to CNNs, while pre-training can mitigate this gap to some extent. Consequently, the relatively stable performance of pre-trained gLMs with limited training data underscores their robustness and efficiency in resource-constrained settings.

Taken together, large gLMs may demonstrate their full potential in more challenging scenarios where long contextual information is available or when training data is limited. Despite their substantial computational resource requirements, these models can capture complex patterns and dependencies that are not easily discernible by simpler models. Conversely, well-designed task-specific methods often excel in handling more straightforward tasks with greater efficiency.

### translation efficiency prediction by mean ribosome loading

Translation regulation is a fundamental biological process that modulates protein synthesis from mRNA, determining when, where and how abundantly proteins are produced within cells. This regulatory mechanism is critical under both normal physiological conditions and in various disease states^54^. The sequence of the 5’ UTR plays a crucial role in regulating mRNA translation efficiency, which can be quantified by the MRL value of mRNA. Despite previous studies identifying a series of regulatory elements within the 5’ UTR regions^54,55^, accurately estimating mRNA translation efficiency remains a significant challenge. To evaluate the capability of gLMs in addressing these questions, we constructed a regression task that required the models to predict the MRL of a given mRNA based on its 5’ UTR sequence.

We employed a large-scale synthetic human 5’UTR library^33^ as the dataset for the MRL task. The training data comprises 76,319 synthetic 5’UTRs, with length ranging from 25 to 100bp, along with their corresponding MRL values. The test data includes 7,600 human 5’UTRs with the same length distribution, and was used to evaluate prediction performance of different models.

Given that this task involves numerical regression, which differs from the classification tasks typically used for pre-training gLMs, we customized the fine-tuning framework for all gLMs^19,24^. Specifically, the embeddings of RNA sequences were extracted from a pre-trained gLM and subsequently input into a downstream network module for the specific task. During this process, only the downstream network was trained (Methods). For comparison, we introduced a CNN-based task-specific method, Optimus^33^. Additionally, we employed the ResNet model as a task-specific method, which is a widely used model in this task and is known for its utilization of skip connections between non-adjacent layers to enable the construction of much deeper convolutional networks^24^.

The results indicated that the fine-tuned gLMs exhibited varied performance scores across multiple evaluation metrics (Fig. 5). Notably, RNAErnie and RNAMSM outperformed all other models according to several criteria, including MSE error (Fig. 5b), MAE error (Fig. 5c), R-squared value(Fig. 5d), Spearman’s correlation coefficient (Fig. 5d), and Pearson’s correlation coefficient (Fig. 5c). Other gLMs maintained competitive scores across these metrics, albeit with slight variations. Interestingly, RNABERT, despite its suboptimal performance in previous tasks, demonstrated competitive results in this task. Both task-specific methods, Optimus and ResNet, did not surpass the pre-trained gLMs but exhibited capabilities comparable to them (Fig. 5e,f). Lastly, DNABERT2 demonstrated worse performance than others.

**Figure 5.**
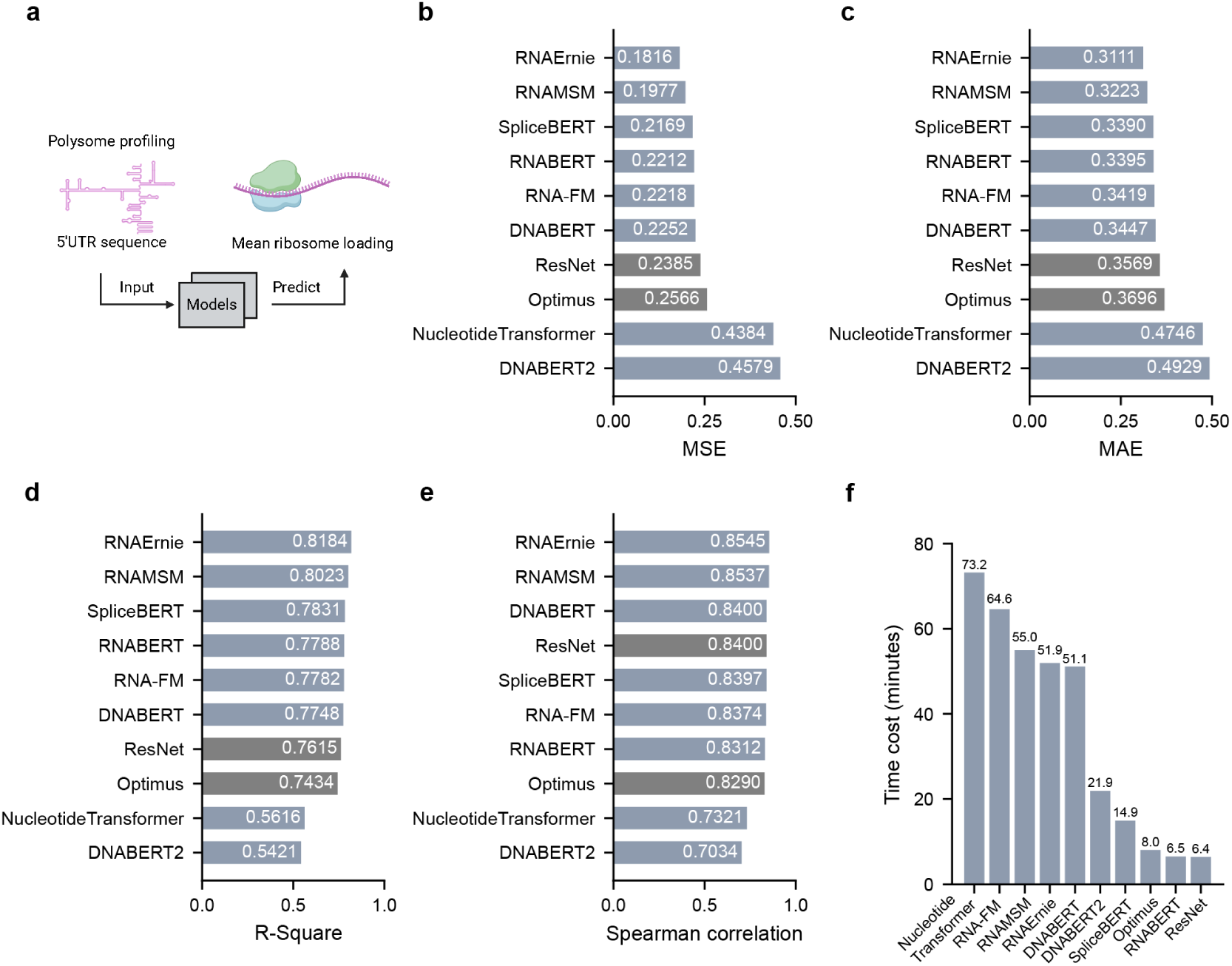
Regression task of predicting mean ribosome loading. **a** Diagram of this task. Multiple measurements comparing across different models: **b** Mean square error, **c**, Mean absolute error **d** R-Square error, and **e** Spearman correlation error. **f** Relative fine-tune time of models for the task.

An interesting observation is that two motif-based methodologies, RNAErnie and DNABERT2, have demonstrated markedly divergent performances in this task. RNAErnie achieved superior results across four out of five evaluation metrics, whereas DNABERT2 did not meet the expected benchmarks in any of the assessments. A key distinction between these models lies in their input processing and training strategies. RNAErnie utilizes nucleotide sequences as input and has undergone specialized pre-training aimed at capturing motif-level characteristics. Conversely, DNABERT2 employs predefined motifs as its input, with the input sequences being fragmented into non-uniform k-mers, where the value of k can extend up to 30bp. As a result, the short input sequence can be represented as one or several tokens. This approach may limit the model’s ability to effectively calculate interactions between different parts of the sequence, as the context provided by nucleotides is reduced. Similarly, the Nucleotide Transformer, which uses a 6-mer encoding that encodes input short 5’UTR sequence into only 4 tokens, might encounter analogous limitations. To serve as a comparative example, the DNABERT model, which underwent pre-training on DNA datasets using standard 3-mers as input, exhibited notably improved performance relative to DNABERT2. These highly specialized motifs may prevent the model from understanding the MRL task in the fine-tuning phase. As a natural control example, the DNABERT model, which was also pre-trained on DNA data but took regular 3-mers as input, gained obviously better performance than DNABERT2. This comparison underscores the notion that a specific input design may not be capable of all general tasks.

What’s more, it can also be observed that the methods RNABERT, ResNet, and Optimus finished the task at a very rapid speed (Supplementary Fig. 6). Again, while the gLMs showed great potential in predicting mRNA translation efficiency, those simpler but specialized methods have also achieved competitive performance at a fraction of the computational cost.

### Proposal of model selection based on resources and biological context

To sum up, we conducted a series of benchmarks on various deep learning models, including eight pre-trained gLMs and multiple task-specific models, applying to prediction tasks covering four RNA biological processes. To highlight the performance differences of the gLMs in different scenarios, we ranked and summarized their results across all evaluations (Fig. 6a). We hope that this ranking provides valuable guidance for researchers looking to apply current pre-trained gLMs to new applications.

**Figure 6.**
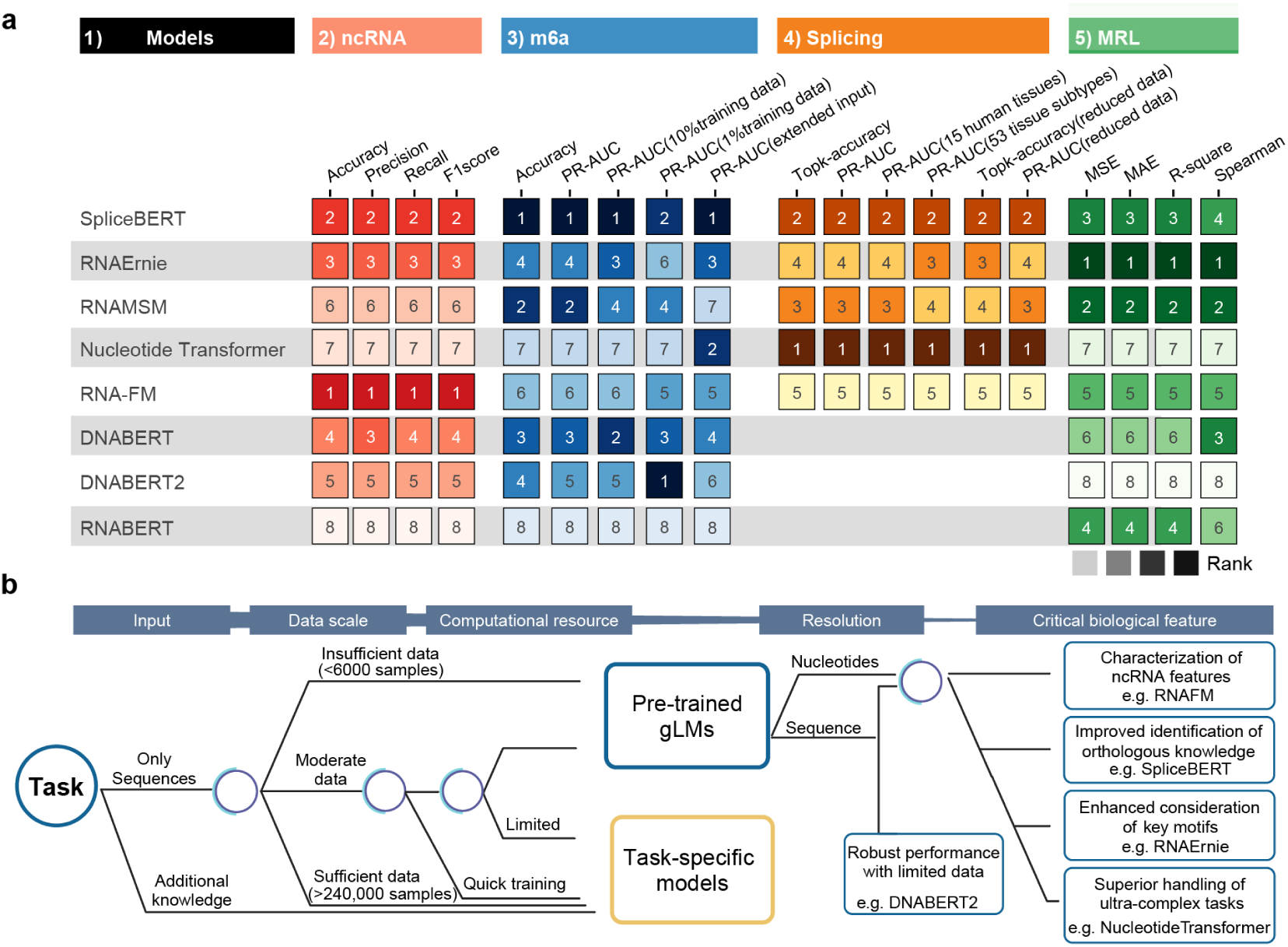
Summary of benchmarking. **a** Ranking of the gLMs across multiple biological tasks. The models were ranked based on their frequency of securing a top-3 position in various tasks. **b** A brief proposal outlines the criteria for selecting deep learning models for de novo tasks.

While gLMs generally excel in performance, the larger model is not always better. Interestingly, models that integrate essential biological features consistently demonstrate superior performance in related tasks. For example, SpliceBERT and Nucleotide Transformer, which incorporated knowledge from multiple vertebrate genomes, performed robustly in tasks of m6a prediction and splicing prediction due to their ability to capture evolutionary conservation. Moreover, Nucleotide Transformer, characterized by its huge training input and training parameters, high model complexity, and long sequence context, exhibited superior performance when processing long input sequences. In contrast, its performance was less impressive among other gLMs in tasks with shorter input sequences. RNAErnie, specifically trained with motif knowledge, showed moderate overall results but performed slightly better in MRL prediction, likely due to its emphasis on RNA binding motifs. RNA-FM, which was particularly pre-trained with a large amount of ncRNA data, captures embeddings of ncRNAs better than other larger models. Nevertheless, this specialization may result in suboptimal performance in other tasks. Notably, the performance differences between these models are relatively small in most cases. While no single model has proved as the unequivocal leader across all tasks, selecting models based on specific biological insights has proven to optimize performance in particular applications.

According to the results in this study, we have designed a recommendation framework for researchers to consider in their RNA-related prediction tasks (Fig. 6b). Several aspects are taken into account in the recommendation framework. Firstly, current pre-trained gLMs are primarily designed to process sequence data as input. For prediction tasks where additional biological information is valuable, task-specific models can more flexibly incorporate such information, thereby solving problems with greater efficiency. Then, based on the ablation tests conducted with varying data sizes, our findings indicate that pre-trained gLMs exhibit robust performance in scenarios with limited training data. Conversely, when sufficient data is available, task-specific methods can achieve comparable performance to gLMs. Empirical evidence from our benchmarks suggests that task-specific methods underperform pre-trained gLMs when training datasets contain less than 6,000 samples. Performance levels between task-specific methods and pre-trained gLMs converge when approximately 50,000 samples are available, and task-specific methods consistently outperform pre-trained gLMs with training datasets exceeding 240,000 samples. It is important to note that these figures should not be considered as absolute thresholds, as variations in task complexity and data quality across different sources can significantly impact these results. Moreover, the balance between time and computational resource efficiency and model performance should also be considered. Our findings indicate that there is no straightforward linear relationship between model performance and time cost. For example, a task-specific method employing a simple CNN achieve approximately 80% performance of gLMs while being 50 times faster in training and inference. Utilizing such a model allows researchers to conduct rapid validation for preliminary analyses, subsequently determining whether a more complex method is warranted for further needs. Additionally, many of these robust gLMs were pre-trained with substantial computational resources, such as 4 to 128 advanced GPUs for training, and 1 to 8 GPUs for fine-tuning. Therefore, it is essential to factor in the computational resource requirements when applying these models.

Our benchmark reveals that when utilizing or developing pre-trained gLMs, the selection should be guided by the critical biological features behind the task. For example, in the task designed to different target resolution in this research, models with motif-based input, such as DNABERT2, exhibit advantages in sequence-level tasks with limited training samples but are not well-suitable for nucleotide-level problems. Other notable aspects of these models include the variations in biological insights obtained from different training datasets and their ability to generalize across different tasks. When faced with tasks focused on ncRNA, RNA-FM may be a better choice due to its training on ncRNA datasets and its advantage in capturing ncRNA features. For complex biological tasks, Nucleotide Transformer, with its ability to process ultra-long input sequences and extract generalizable pattern, is advantageous. When dealing with motif-related problems, RNAErnie, with its motif-focused training, can be particularly effective. In RNA splicing related tasks, SpliceBERT, leveraging conserved sequences and structures across multiple vertebrate genomes, demonstrates strong performance. These considerations underscore the importance of selecting the appropriate model based on specific biological insights and task requirements.

These types of models represent various directions explored by researchers in the field and may exhibit flexibility in different scenarios. According to our benchmarking analysis, no current gLM demonstrates a dominant advantage across all biological tasks. However, we anticipate that future large-scale biological language models will integrate these critical aspects. By doing so, a more powerful biological tool may emerge, significantly advancing the evolution of bioinformatics research.

## Discussion

Accurately predicting RNA-related biological events from sequence data has historically presented significant challenges. However, recent advances in computational methodologies, particularly the development of large biological deep models, are transforming this landscape. Inspired by the monumental achievements of language models in Natural Language Processing (NLP), researchers have introduced a series of pre-trained Genomic Language Models. These models leverage self-supervised training on extensive genomic datasets to capture intrinsic features of RNA sequences. Subsequently, they can be fine-tuned for diverse biological tasks, demonstrating their potential to revolutionize RNA biology research.

While pre-trained gLMs hold significant promise, it remains an open question whether they can effectively manage all RNA-related tasks and outperform algorithms tailored for specific tasks. In this study, we benchmarked the performance of eight pre-trained gLMs alongside several task-specific methods across four representative biological tasks. Our evaluation extends beyond mere performance comparison; we also delve into the particular contexts in which these models exhibit substantial advantages.

Our findings indicate that pre-trained gLMs outperform task-specific methods in various scenarios, particularly when data is scarce for certain biological tasks. Despite extensive evaluations across numerous tasks, the extent to which the gLMs can hold multiple de novo inquiries remains uncertain. The strength of these models derives from distinct design strategies. For instance, RNAErnie leverages its ERNIE training framework to enhance motif-related sequence feature learning; SpliceBERT excels in capturing evolutionary conservation features by pre-training on genomics across different species, giving it an edge in several tasks; RNA-FM, though lacking extensive special designs, benefits from a large training dataset and prolonged pre-training, resulting in consistent, moderate performance across tasks; and Nucleotide Transformer integrates ultra-long sequence features to tackle highly complex tasks especially those context related problems.

Task-specific models, though typically trained with fewer resources than pre-trained gLMs, can sometimes achieve competitive performance by holding unique properties not present in larger gLMs. For instance, ncRDense incorporates RNA structural information as additional input for the ncRNA classification task; SpTransformer and SpliceAI utilize extra context from thousands of flanking nucleotides in splicing prediction tasks. These strategies enable them to match or surpass gLMs, but cannot easily replicate by current pre-trained models due to algorithmic or hardware constraints. Furthermore, our comparative analysis highlights that as available training data increases, the advantage of pre-trained gLMs diminishes. Additionally, lightweight task-specific models offer the benefit of rapid training and application, making them valuable in certain research contexts requring time efficiency.

When faced with a new biological challenge, selecting the most suitable methodology can be daunting. Our preliminary findings offer guidance to assist researchers in making informed choices. Although this guidance may not be universally applicable, we hope it will prove valuable in future investigative efforts and contribute new insights to the field.

Despite pre-trained gLMs demonstrating reliable performance across tested scenarios, no single model consistently outperforms all others in every situation. This study highlights the current limitations of various gLMs in biological applications. However, these challenges are not insurmountable and can be addressed through improvements in algorithmic details. Looking ahead, we anticipate continued advancements in AI technology within the realm of biology. By overcoming present limitations, we believe that large models will provide researchers with more powerful tools, thereby propelling biological research and fostering breakthroughs in healthcare, biotechnology, and other related fields.

## Methods

### Overview of the benchmark

To advance the development of RNA studies, we have established a unified benchmarking framework in providing reasonable comparisons and meaningful insights into the development of deep learning algorithms for RNA sequence-related biological tasks. Four types of tasks were considered for benchmarking: 1) Multi-class classification for the whole sequence; 2) Classification for a single position; 3) Classification for each nucleotide; 4) Regression for sequences. We selected specific biological tasks for each type of task to investigate whether current gLMs could handle all types of downstream tasks. The code framework is adaptable, allowing for the inclusion of additional tasks for evaluation. In total, eight open-source gLMs were included in our benchmark: RNA-FM^24^, RNAMSM^25^, RNABERT^23^, SpliceBERT^20^, DNABERT^27^, DNABERT2^21^, and RNAErnie^19^. We obtained pre-trained model weights from each respective publication and fine-tuned them for downstream tasks. In particular, the official implementation of RNAErnie was based on the PaddlePaddle Machine Learning Framework, which was not directly compatible with the Pytorch framework that was used by all other gLMs. Therefore, we incorporated the Pytorch version of RNAErnie recommended by the authors (see Code Availability section).

### Evaluation metrics

We utilized multiple metrics for the tasks. The definitions of these metrics are as follows:

- Area Under the Receiver Operating Characteristic Curve (ROC-AUC): ROC-AUC measures the model’s ability to classify samples at different thresholds. It is calculated as the area under the receiver operating characteristic curve, which is plotted as a curve with the false positive rate on the x-axis and the true positive rate on the y-axis. The ROC-AUC value ranges from 0 to 1, with 1 indicating perfect classification and 0.5 representing random guessing.
- Precision: It is the ratio of true positive observations to the total predicted positives. It indicates the accuracy of the positive class predictions.

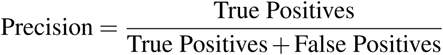
- Recall: Also called sensitivity, is the ratio of true positive observations to the actual positives. It measures the ability of a model to identify all relevant instances.

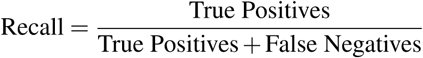
- Area Under the Precision-Recall Curve (PR-AUC): PR-AUC is a performance metric used to evaluate binary classifiers, which measures the average precision across different recall levels. It can be used to evaluate the performance of classifiers in the presence of imbalanced classes or uneven sample distributions. PR-AUC is a more sensitive metric than ROC-AUC, particularly for the classification of imbalanced data.
- F1-score: The F1-score is the harmonic mean of precision and recall, providing a balance between the two metrics. It is especially useful when the class distribution is uneven.

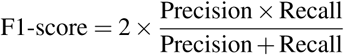
- Accuracy: Accuracy is the ratio of correctly predicted observations to the total observations. While useful, it can be misleading for imbalanced datasets.

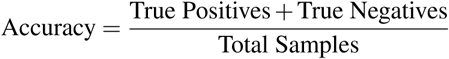
- Top-k accuracy: Top-k accuracy measures the fraction of times the true label is among the top k predicted labels.
- Mean Squared Error (MSE): MSE is a common metric for regression tasks, representing the average of the squares of the errors between predicted and actual values. It penalizes larger errors more than smaller ones.

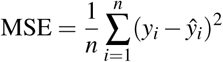
- Mean Absolute Error (MAE): MAE measures the average magnitude of the errors in a set of predictions, without considering their direction. It is the average over the absolute differences between predicted and actual values.

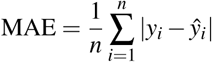
- R-square (R-squared coefficient of determination): R-square indicates how well the independent variables explain the variability of the dependent variable in a regression model. It ranges from 0 to 1, with higher values indicating better model fit.

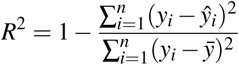
- Spearman correlation: It assesses how well the relationship between two variables can be described using a monotonic function. It is a non-parametric measure of rank correlation.

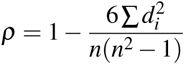

where *d_i_* is the difference between the ranks of each observation, and *n* is the number of samples.

### Standardized Fine-tuning gLMs for RNA biological tasks

To ensure fair evaluation across various methods and tasks, we implemented general and standardized training and testing pipelines. detailed descriptions for each section are provided below. Given that most gLMs had not previously encountered these specific tasks, we selected hyperparameters based on those used in related tasks by the models. We aimed to maintain consistent training settings across all models for the same tasks, with exceptions only where necessary (e.g., models requiring excessive GPU resources might have reduced batch sizes).

During training and testing, all models processed the same RNA sequences as input, despite the nucleotide ‘U’ replaced by ‘T’ in some models. To ensure fair evaluation, all models were trained for an equal number of epochs. The best epoch was determined based on specific metrics (detailed in the Methods section for each task), and the performance of this best epoch was reported in the figures. Pre-trained gLMs required input sequences to be annotated with special tokens like “[CLS]”, “[SEP]”, or “[PAD]”, which varied among models. Therefore, references to “512bp input” imply an actual valid input length of 511bp or 510bp. These details are documented in our source code.

### Construction of ncRNA classification task

The non-coding RNA (ncRNA) classification task involves models categorizing input ncRNA sequences into one of 13 predefined categories. The dataset was constructed based on the Rfam database release 12^29,39^, consisting of 6,320 samples in the training set and 2,600 samples in the test set. Each input sequence was assigned a label *y* ∈ 0, 1*, …,* 12 representing its class, which includes categories such as miRNA, 5S rRNA, 5.8S rRNA, ribozymes, CD-box, HACA-box, scaRNA, tRNA, Intron gpI, Intron gpII, IRES, leader, and riboswitch.

For each gLM, a convolution layer was appended to the backbone network to derive the output score from sequence embeddings provided by the pre-trained backbone networks. The models were trained using parameters listed in Supplementary Table. 1, employing the AdamW optimizer and Cross-Entropy loss function. After each training epoch, models were evaluated using multiple metrics. The baseline values of the task-specific model ncRDense were sourced from cited literature. We did not retrain ncRDense as the literature provided online tools but no official source code. Performance metrics for nRC and RNACon were also taken from the same publication.

The evaluation employed Accuracy, Precision, Recall, and F1-score metrics. For each model, the epoch with the highest F1-score value was identified as the best epoch, and other metrics from this epoch were reported. Model performance across all epochs is presented in Supplementary Fig. 1. Results of nRC and RNACon were directly cited from corresponding respective publications.

### Construction of m6A prediction task

The m6A prediction task requires models to predict whether the center of an input sequence undergoes m6a modification. The training dataset comprises 24,518 positive samples and 24,532 negative samples, while the test dataset includes 6,304 positive samples and 6,307 negative samples. Each input sequence is 101 bp in length, with 50 bp flanking sequences upstream and downstream of the center, and is assigned a binary label *y* ∈ 0, 1. An extra dataset with extended input length was generated, where sequences were extended to 509 bp (254 bp flanking sequences for each side, considering that most gLMs accept a maximum input length of 512 bp. Special tokens “CLS” and “SEP” were added to the input sequences as required by the models. Similar to the ncRNA task, a convolution layer was appended to the backbone network of each gLM to derive output scores from sequence embeddings provided by the pre-trained backbone networks. The task-specific deep model, DeepM6ASeq, was included in the pipeline without modification. Models were trained using parameters listed in Supplementary Table. 1, employing the AdamW optimizer and Binary Cross-Entropy loss function. For each model, the epoch with the highest PR-AUC value was identified as the best epoch, and other metrics of this epoch were reported. Model performance across all epochs is displayed in Supplementary Fig. 2.

### Construction of tissue-specific alternative splicing prediction task

The splicing prediction task requires models to distinguish all splice sites among the center 500bp of an input sequence. The datasets were constructed from the GTEx V8^53^ database,, which consists of 838 donors, providing 17,382 samples from 53 tissues and two cell lines. To obtain meaningful splice sites and the corresponding tissue usage ratios, we processed the exon-exon junction read counts file from the dataset for each tissue type. To compile the datasets, pre-mRNA sequences of each gene were extracted based on the GRCh38 reference genome. Splice junctions from the GTEx database were considered, defining the bases preceding and following each splice junction (i.e., the 5’ and 3’ ends of exons) as splice sites. For genes with multiple transcripts, sequences were extracted from the most upstream site across all transcripts to the most downstream site. Each sequence was then divided into 500 bp blocks, discarding any that did not contain splice sites.

Splice site positions were labeled as “acceptor” or “donor” if supported by any sample without conflict. The exon start site splice sites were labeled “acceptor,” and the end sites were labeled “donor,” with all other positions labeled “neither.” The tissue usage label for each splice site represented the proportion of samples from the tissue containing corresponding splice junctions:

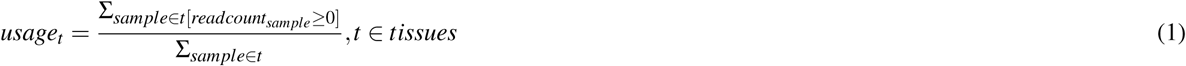

For each remaining block, the flanking sequence and corresponding 500 bp label were packaged as a single training (testing) data entry. The length of the flanking sequence was determined by model limitations. During training and testing, any splice site with a maximum usage label below 0.05 across all tissue classes was re-labeled as “neither”. Three detailed tasks were considered: 1) Distinguish “acceptor”, “donor” and “neither” for each nucleotide. 2) Predict tissue usage for 15 human tissues. 3) Predict tissue usage for 53 detailed tissue subtypes. Sequences from chromosomes 2, 4, 6, 8, 10-22, X, and Y were used to create the training dataset, while sequences from chromosomes 1, 3, 5, 7, and 9 formed the testing dataset. We ensured no test data was exposed in the training data by cross-referencing sequence strands with the Ensembl database (release 110) and excluding gene sequences with paralogs. Despite independent dataset splitting, we verified no overlap between training and testing data. The resulting dataset contains 242,017 training entries (121,008,500 labeled nucleotides) and 99,813 testing entries (49,906,500 labeled nucleotides). For each model, the epoch with the highest average PR-AUC value across all classes was recognized as the best epoch, with other metrics presented accordingly.

As an ablation test, we constructed a lightweight dataset, which includes chromosomes 2, 4, 6, 8, 10 as the training dataset, and chromosome 1, 3, 5, 7, 9 as the testing dataset. Other details were the same as the above description.

Two metrics, Top-k accuracy and PR-AUC, were used for evaluation. For multi-label classification (15 classes and 53 classes), performance was calculated for each class, and the mean PR-AUC values were used for ranking the models.

### Construction of translation efficiency prediction task

The translation efficiency prediction task requires models to evaluate the numerical mean ribosome loading value of transcripts based on the input 5’ UTR sequence. The dataset was derived from a synthetic human 5’UTR library^33^. The training dataset consists of 83,919 5’UTRs of 75 different lengths ranging from 25 bp to 100 bp and their corresponding MRLs, while the testing dataset includes 7,600 real human 5’UTRs with the same length distribution provided by the same library. MRL labels were standardized based on the distribution of the training data. Since all the models were pre-trained on classification datasets and lacked prior exposure to value regression, fine-tuning the backbone network was not suitable^19,24^. Following the RNA-FM pipeline for handling numerical regression tasks, we froze the backbones of gLMs. A ResNet structure was appended to the backbone model instead of a simple convolution layer. This ResNet takes the gLMs’ embeddings as input and predicts a value for the input sequence, with the backbone model weights remaining unchanged during training. Adam optimizer and Binary Cross-Entropy (BCE) Loss function were employed. For each model, the epoch with the best Mean Square Error (MSE) value was identified as the best epoch, with other metrics from this epoch presented. As an ablation test, a bare ResNet model was included in the comparison, which uses one-hot encoded sequences instead of sequence embeddings as input. All models employed the AdamW optimizer and BCE Loss function.

The evaluation employed metrics including MSE, Mean Absolute Error (MAE), R-square, and Spearman correlation.

## Data availability

All public data and tools utilized in the analysis are referenced in the main text or the supplementary materials. The utilized public datasets are listed below: The Rfam database was published in^39^. The related m6A dataset was published in^10^. GTEx exon-exon junctions data are available from the GTEx Analysis V8 (dbGaP Accession phs000424.v8.p2)[https://www.ncbi.nlm.nih.gov/projects/gap/cgi-bin/study.cgi?study_id=phs000424.v8.p2]; The hg38 genome annotation and gene paralogs data [www.ensembl.org]; The mean ribosome loading data was published in^33^.

## Code availability

Code about the benchmark framework and analysis pipeline is available under an Apache-2.0 license on GitHub (https://github.com/ShenLab-Genomics/biombenchmark).

## Acknowledgements

We thank the technical support by the Core Facilities of Liangzhu Laboratory. This work was supported by the National Natural Science Foundation of China (82471904) and fundings from Liangzhu Laboratory. A subset of images (Figures 1, 2a, 3a, 4a, 5a, and 6b) were created with BioRender.com, with permission.

## Author contributions statement

N.Y. designed and developed the benchmark pipeline, collected data, performed data analysis, and drafted the original manuscript; C.L. drafted the original manuscript and revised the manuscript. H.L. contributed to code development and performed data analysis; S.W. supervised the work and contributed to the manuscript; G.C. supervised the work and contributed to the manuscript; N.S. conceived, designed, guided and supervised the study, and revised the manuscript. All authors read and approved the manuscript.

## Competing interests

The authors declare that they do not have any competing interests.

---

**Supplementary Table 1:**
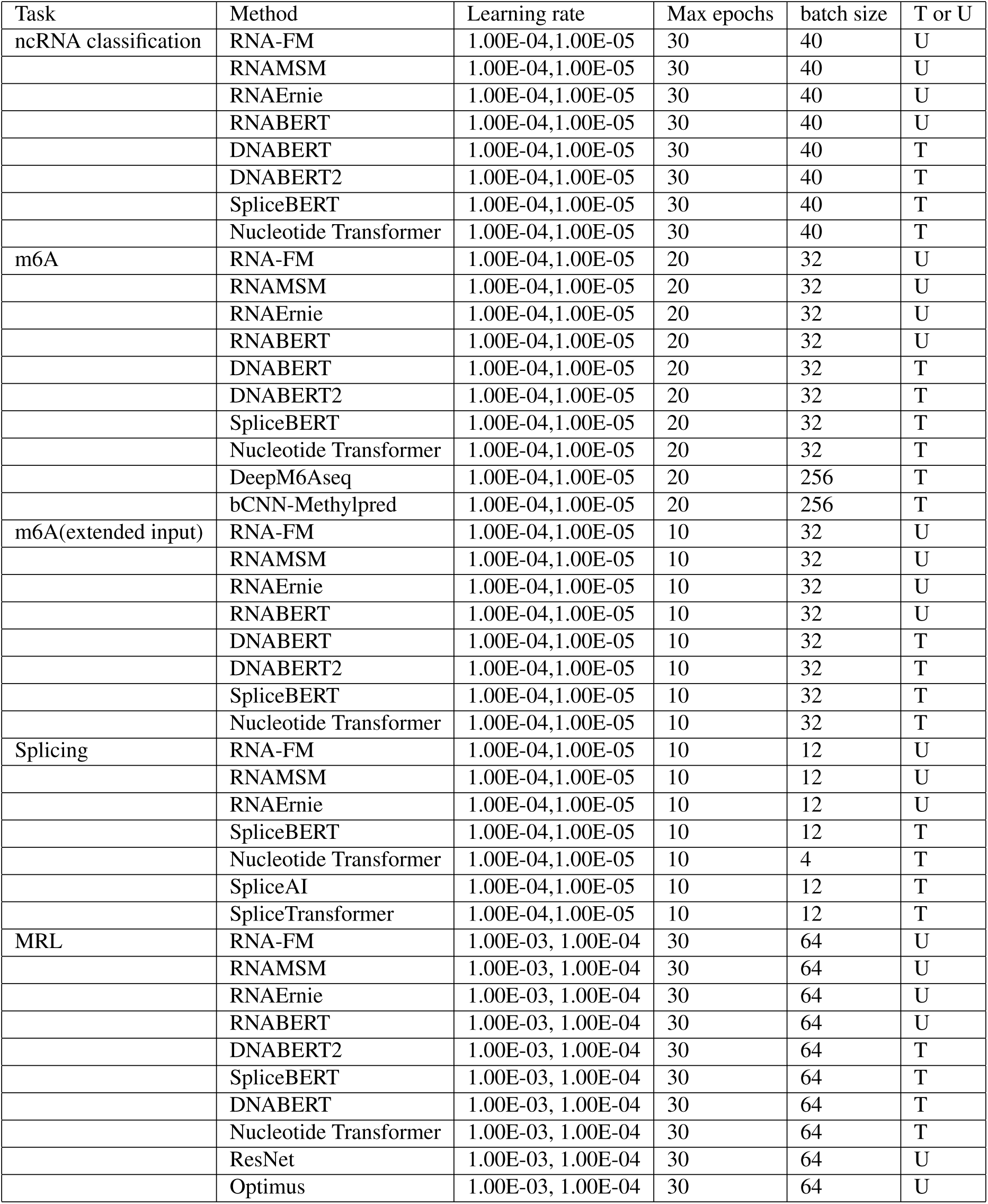
Parameters in training or fine-tuning.

**Supplementary Figure 1:**
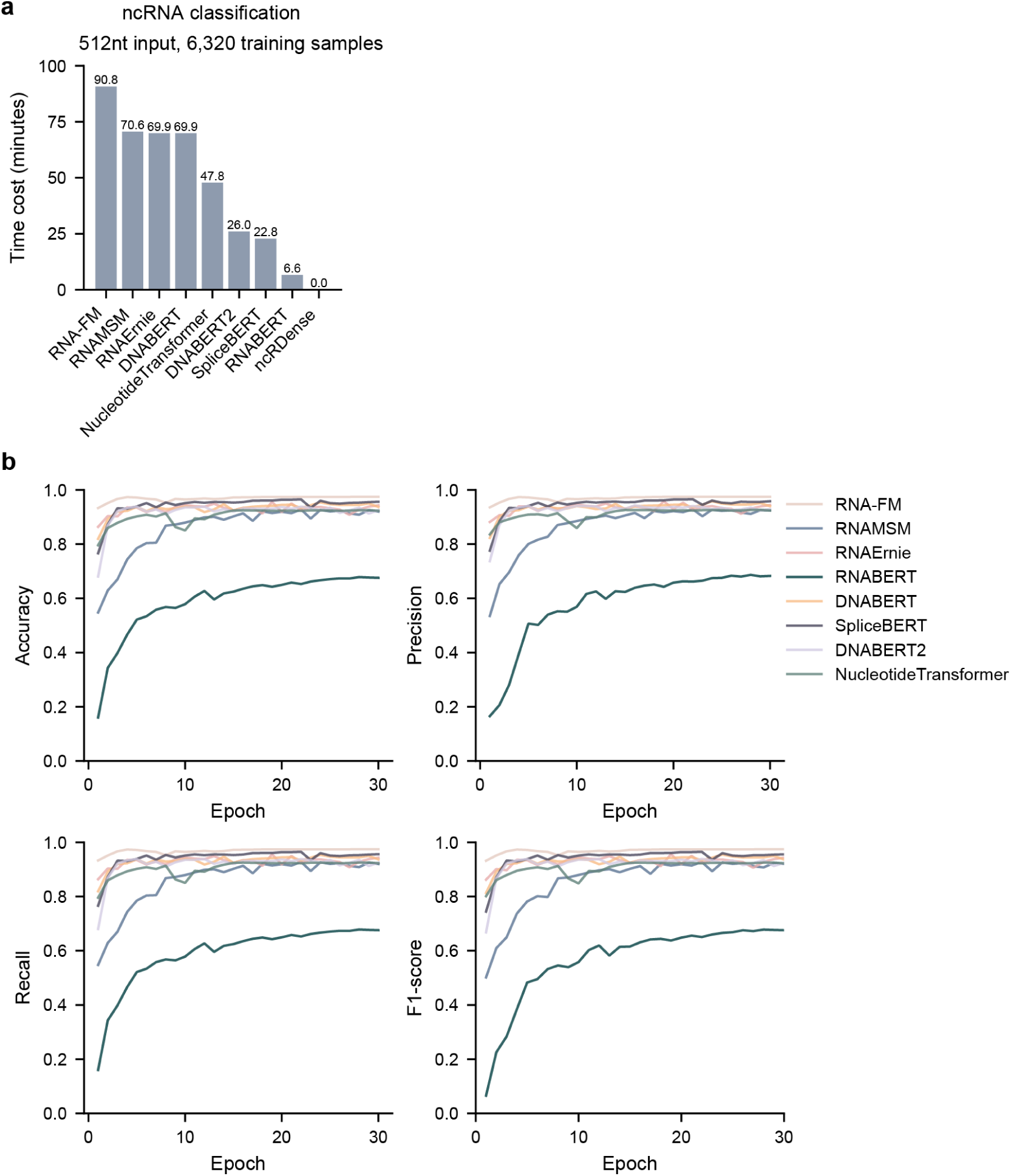
Properties of pre-trained gLMs in the ncRNA classification task. a Average time cost for each epoch in fine-tuning the models. b Performance curves in fine-tuning the models.

**Supplementary Figure 2:**
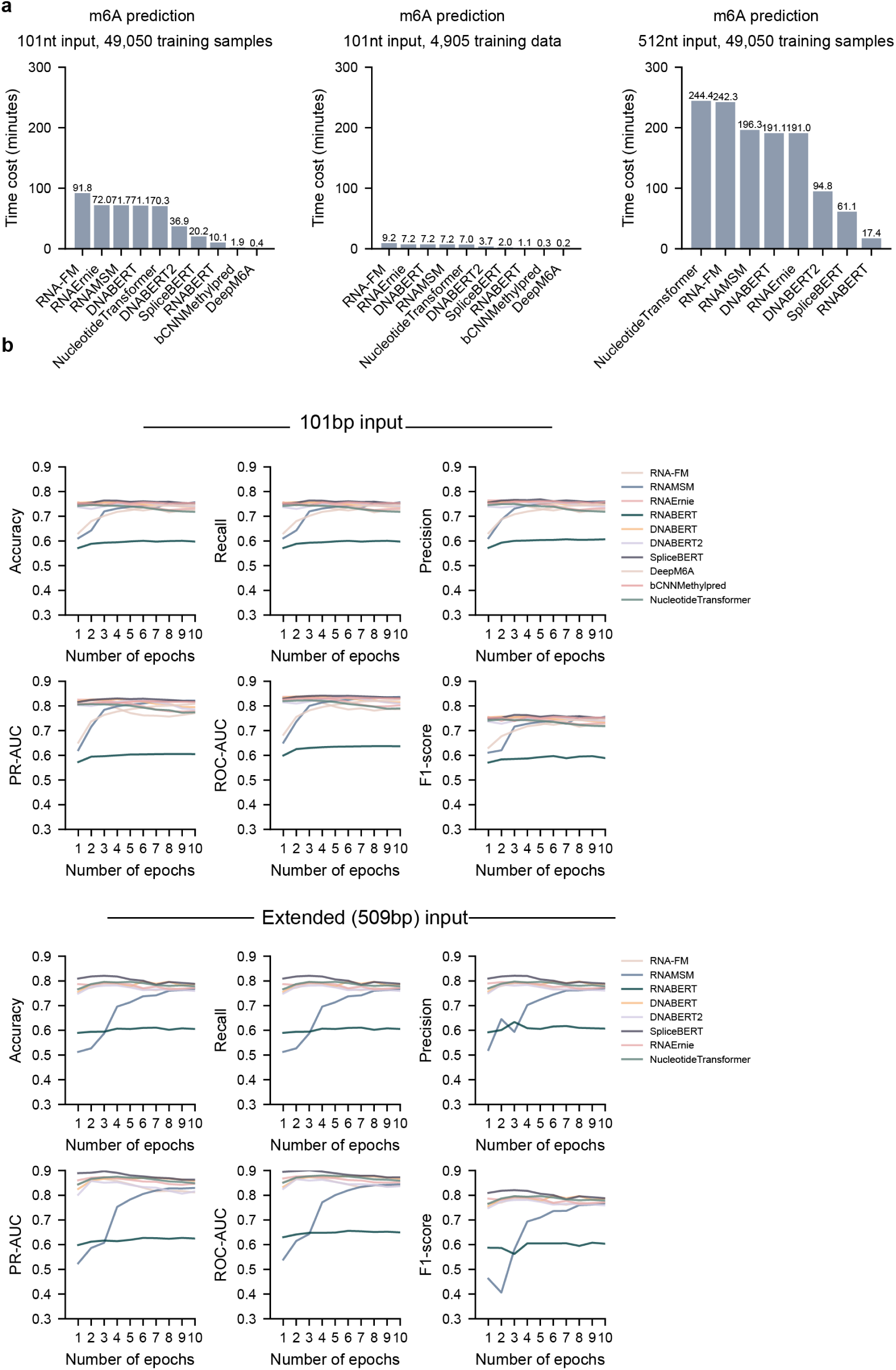
Properties of pre-trained gLMs in the m6A prediction task. a Average time cost for each epoch in fine-tuning the models. b Performance curves in fine-tuning the models.

**Supplementary Figure 3:**
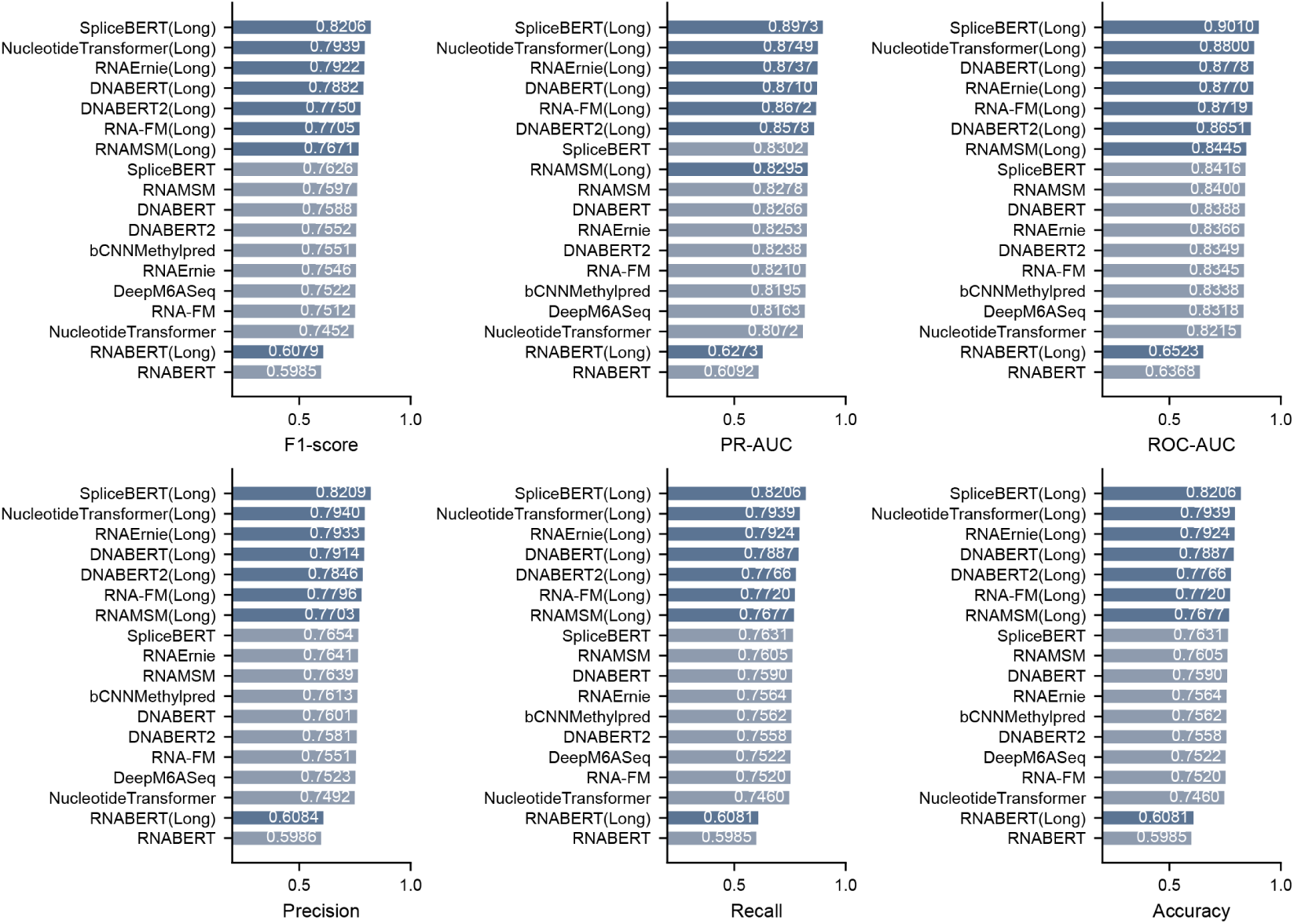
Comparison of model performance using original input vs. extended input (Labeled as ‘Long’).

**Supplementary Figure 4:**
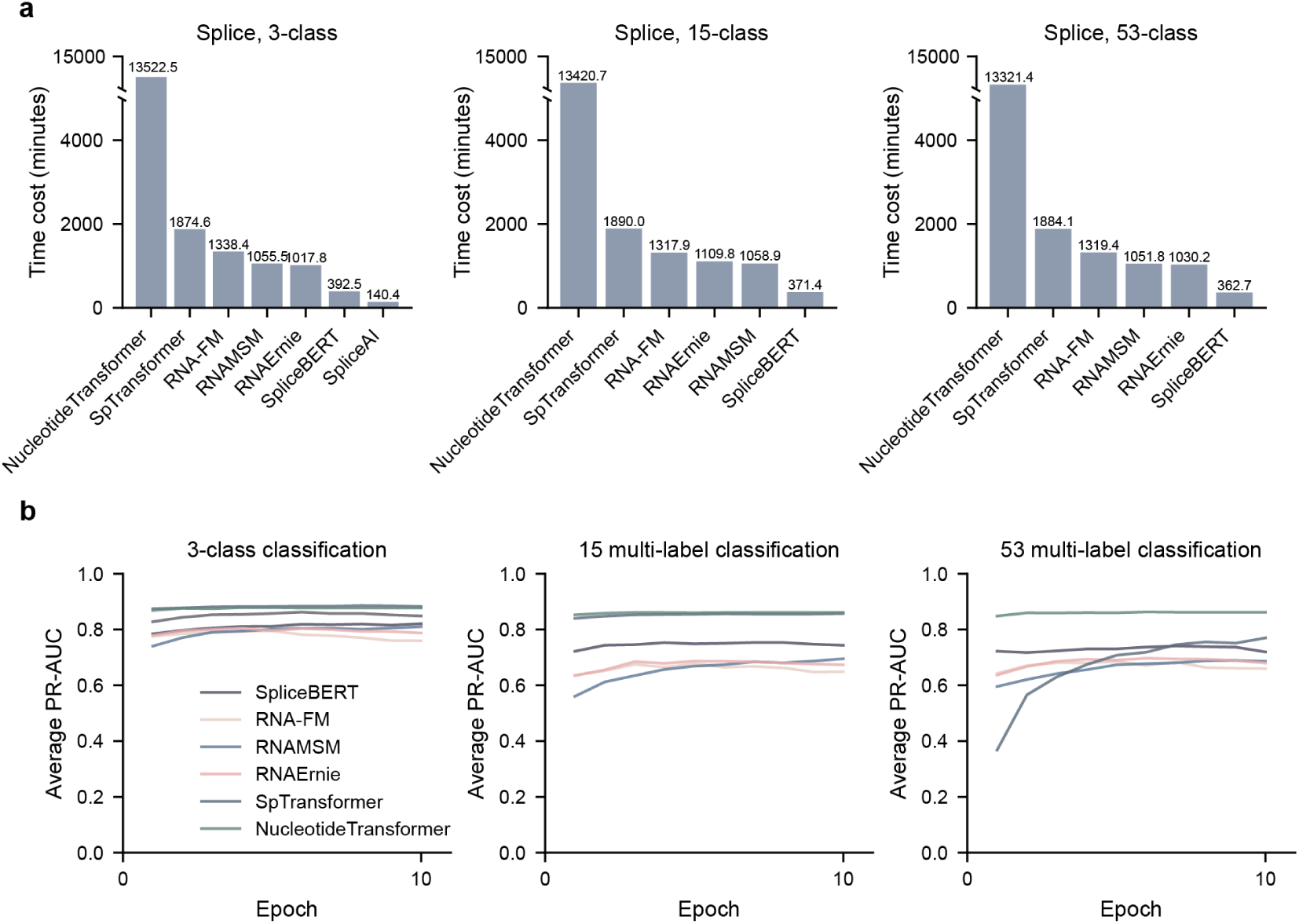
Properties of pre-trained gLMs in the splicing prediction task. a Average time cost for each epoch in fine-tuning the models. b Performance curves in fine-tuning the models.

**Supplementary Figure 5:**
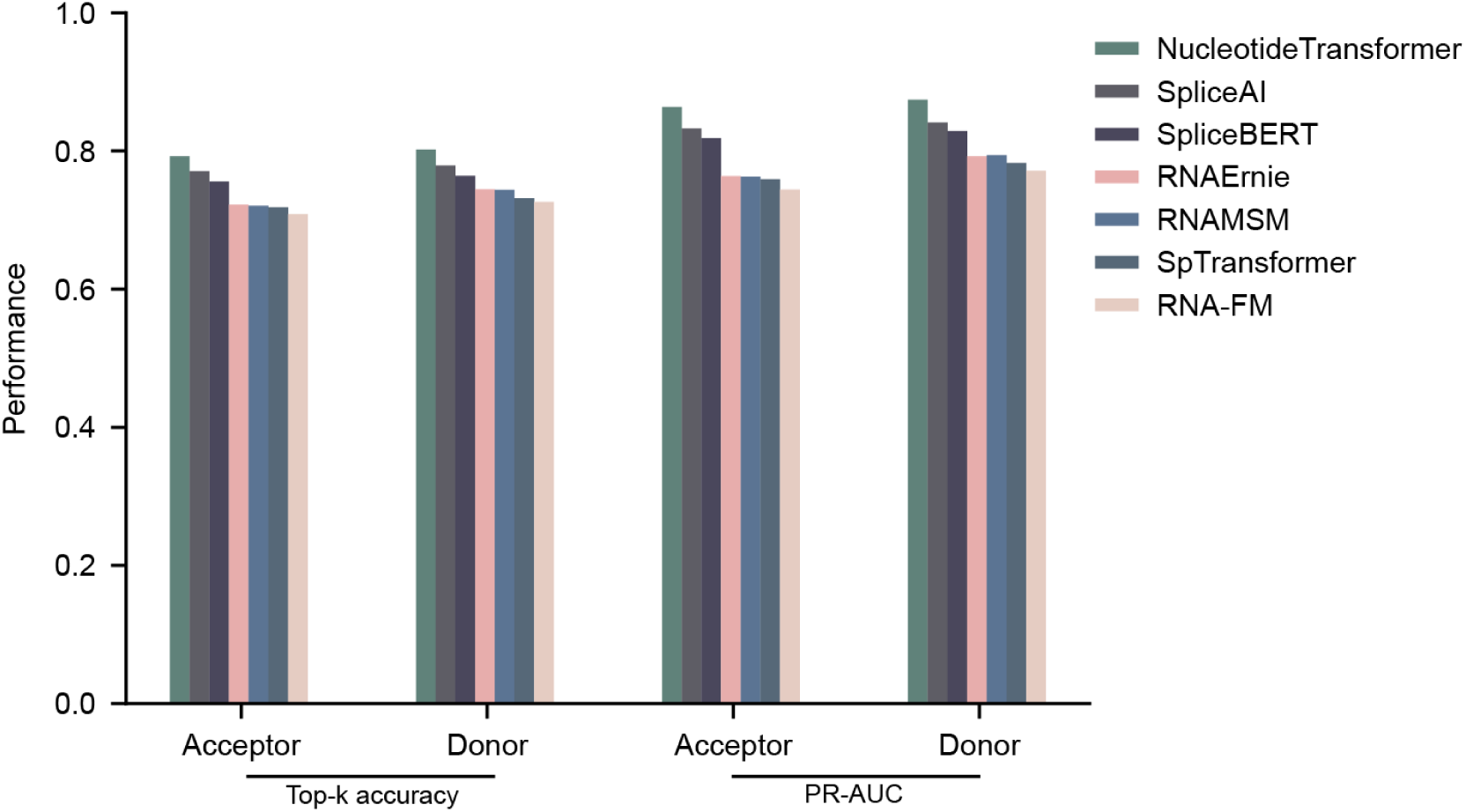
Model performance with a smaller training dataset derived from 5 chromosomes.

**Supplementary Figure 6:**
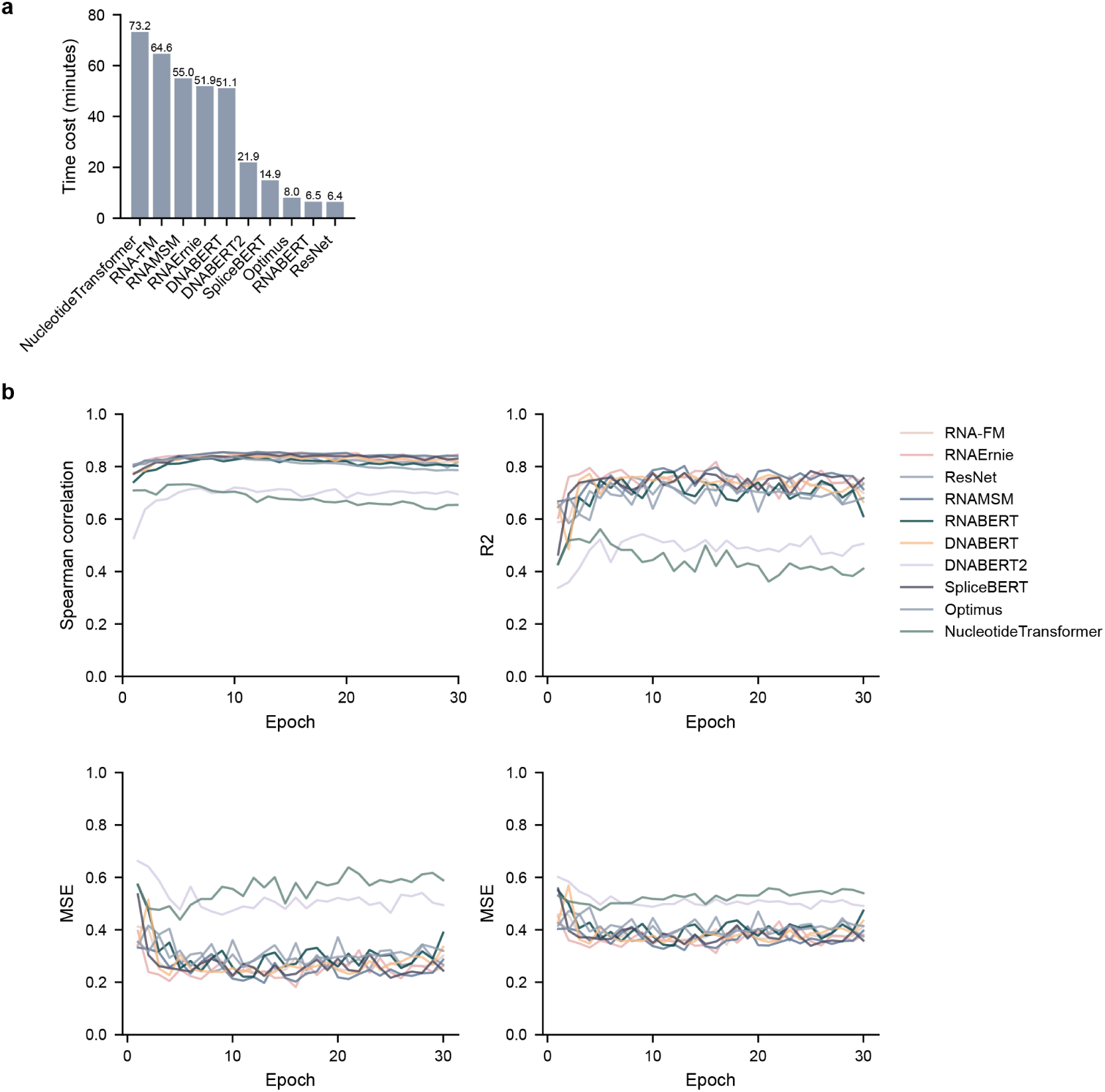
Properties of pre-trained gLMs in the MRL prediction task. a Average time cost for each epoch in fine-tuning the models. b Performance curves in fine-tuning the models.

## Notes

### Competing Interest Statement

The authors have declared no competing interest.

## References

1 Caprara, M. & Nilsen, T. Rna: Versatility in form and function. Nat. structural biology 7, 831–3, DOI: 10.1038/82816 (2000).

2 Emde, A. et al. Dysregulated mirna biogenesis downstream of cellular stress and als-causing mutations: a new mechanism for als. The EMBO J. 34, 2633–2651, DOI: 10.15252/embj.201490493 (2015).

3 Li, Y. et al. Globally reduced n6-methyladenosine (m6a) in c9orf72-als/ftd dysregulates rna metabolism and contributes to neurodegeneration. Nat. Neurosci. 26, 1328–1338, DOI: 10.1038/s41593-023-01374-9 (2023).

4 Humphrey, J., Emmett, W., Fratta, P., Isaacs, A. M. & Plagnol, V. Quantitative analysis of cryptic splicing associated with tdp-43 depletion. BMC medical genomics 10, 1–17 (2017).

5 Breaker, R. Natural and engineered nucleic acids as tools to explore biology. Nature 432, 838–45, DOI: 10.1038/nature03195 (2005).

6 Wang, R., Helbig, I., Edmondson, A. C., Lin, L. & Xing, Y. Splicing defects in rare diseases: transcriptomics and machine learning strategies towards genetic diagnosis. Briefings Bioinforma. 24, bbad284 (2023).

7 Carracedo-Reboredo, P. et al. A review on machine learning approaches and trends in drug discovery. Comput. structural biotechnology journal 19, 4538–4558 (2021).

8 Alharbi, F. & Vakanski, A. Machine learning methods for cancer classification using gene expression data: A review. Bioengineering 10, 173 (2023).

9 Soleymani, F., Paquet, E., Viktor, H., Michalowski, W. & Spinello, D. Protein–protein interaction prediction with deep learning: A comprehensive review. Comput. Struct. Biotechnol. J. 20, 5316–5341 (2022).

10 Zhang, Y. & Hamada, M. Deepm6aseq: prediction and characterization of m6a-containing sequences using deep learning. BMC bioinformatics 19, 1–11 (2018).

11 Jaganathan, K. et al. Predicting splicing from primary sequence with deep learning. Cell 176, 535–548 (2019).

12 You, N. et al. SpliceTransformer predicts tissue-specific splicing linked to human diseases. Nat. Commun. 15, 9129, DOI: 10.1038/s41467-024-53088-6 (2024).

13 Mantegna, R. N. et al. Linguistic features of noncoding dna sequences. Phys. review letters 73, 3169 (1994).

14 Abramson, J. et al. Accurate structure prediction of biomolecular interactions with alphafold 3. Nature 1–3 (2024).

15 Lin, Z. et al. Evolutionary-scale prediction of atomic-level protein structure with a language model. Science 379, 1123–1130 (2023).

16 Benegas, G., Ye, C., Albors, C., Li, J. C. & Song, Y. S. Genomic language models: Opportunities and challenges. arXiv preprint arXiv:2407.11435 (2024).

17 Simon, E., Swanson, K. & Zou, J. Language models for biological research: a primer. Nat. Methods 21, 1422–1429 (2024).

18 Devlin, J. Bert: Pre-training of deep bidirectional transformers for language understanding. arXiv preprint arXiv:1810.04805 (2018).

19 Wang, N. et al. Multi-purpose rna language modelling with motif-aware pretraining and type-guided fine-tuning. *Nat*. Mach. Intell. 6, 1–10, DOI: 10.1038/s42256-024-00836-4 (2024).

20 Chen, K. et al. Self-supervised learning on millions of primary rna sequences from 72 vertebrates improves sequence-based rna splicing prediction. Briefings Bioinforma. 25, DOI: 10.1093/bib/bbae163 (2024).

21 Zhou, Z. et al. Dnabert-2: Efficient foundation model and benchmark for multi-species genome (2023). 2306.15006.

22 Dalla-Torre, H. et al. Nucleotide transformer: building and evaluating robust foundation models for human genomics. Nat. Methods 1–11 (2024).

23 Akiyama, M. & Sakakibara, Y. Informative RNA base embedding for RNA structural alignment and clustering by deep representation learning. NAR Genomics Bioinforma. 4, lqac012, DOI: 10.1093/nargab/lqac012 (2022). https://academic.oup.com/nargab/article-pdf/4/1/lqac012/42577169/lqac012_supplemental_file.pdf.

24. Chen, J., et al. Interpretable rna foundation model from unannotated data for highly accurate rna structure and function predictions. arXiv preprint arXiv:2204.00300 (2022).

25 Zhang, Y. et al. Multiple sequence alignment-based rna language model and its application to structural inference. Nucleic acids research 52, DOI: 10.1093/nar/gkad1031 (2023).

26. Rao, R. M. et al. Msa transformer. In Meila, M. & Zhang, T. (eds.) Proceedings of the 38th International Conference on Machine Learning, vol. 139 of Proceedings of Machine Learning Research, 8844–8856 (PMLR, 2021).

27 Ji, Y., Zhou, Z., Liu, H. & Davuluri, R. V. DNABERT: pre-trained Bidirectional Encoder Representations from Transformers model for DNA-language in genome. Bioinformatics 37, 2112–2120, DOI: 10.1093/bioinformatics/btab083 (2021). https://academic.oup.com/bioinformatics/article-pdf/37/15/2112/50578892/btab083.pdf.

28 Chantsalnyam, T., Siraj, A., Tayara, H. & Chong, K. T. ncrdense: a novel computational approach for classification of non-coding rna family by deep learning. Genomics 113, 3030–3038 (2021).

29 Fiannaca, A., La Rosa, M., La Paglia, L., Rizzo, R. & Urso, A. nrc: non-coding rna classifier based on structural features. BioData mining 10, 1–18 (2017).

30 Panwar, B., Arora, A. & Raghava, G. P. Prediction and classification of ncrnas using structural information. BMC genomics 15, 1–13 (2014).

31 Islam, N. U. & Park, J. bcnn-methylpred: Feature-based prediction of rna sequence modification using branch convolutional neural network. Genes 12 (2021).

32 Koonce, B. & Koonce, B. Resnet 50. Convolutional neural networks with swift for tensorflow: image recognition dataset categorization 63–72 (2021).

33 Sample, P. J. et al. Human 5’ utr design and variant effect prediction from a massively parallel translation assay. *Nat*. biotechnology 37, 803–809 (2019).

34 Nemeth, K., Bayraktar, R., Ferracin, M. & Calin, G. A. Non-coding rnas in disease: from mechanisms to therapeutics. Nat. Rev. Genet. 25, 211–232 (2024).

35 Esteller, M. Non-coding rnas in human disease. Nat. reviews genetics 12, 861–874 (2011).

36 Mattick, J. S. The genetic signatures of noncoding rnas. PLoS genetics 5, e1000459 (2009).

37 Ashrafizadeh, M. et al. Non-coding rna-based regulation of inflammation. In Seminars in immunology, vol. 59, 101606 (Elsevier, 2022).

38 Amaral, P. et al. The status of the human gene catalogue. Nature 622, 41–47 (2023).

39 Nawrocki, E. P. et al. Rfam 12.0: updates to the rna families database. Nucleic acids research 43, D130–D137 (2015).

40 Batista, P. J. et al. m6a rna modification controls cell fate transition in mammalian embryonic stem cells. Cell stem cell 15, 707–719 (2014).

41 Berlivet, S., Scutenaire, J., Deragon, J.-M. & Bousquet-Antonelli, C. Readers of the m6a epitranscriptomic code. Biochimica et Biophys. Acta (BBA)-Gene Regul. Mech. 1862, 329–342 (2019).

42 Meyer, K. D. et al. Comprehensive analysis of mrna methylation reveals enrichment in 3’ utrs and near stop codons. Cell 149, 1635–1646 (2012).

43 Linder, B. et al. Single-nucleotide-resolution mapping of m6a and m6am throughout the transcriptome. Nat. methods 12, 767–772 (2015).

44 Wang, Z. & Burge, C. Splicing regulation: From a parts list of regulatory elements to an integrated splicing code. RNA (New York, N.Y.) 14, 802–13, DOI: 10.1261/rna.876308 (2008).

45 Tazi, J., Bakkour, N. & Stamm, S. Alternative splicing and disease. Biochimica et Biophys. Acta (BBA)-Molecular Basis Dis. 1792, 14–26 (2009).

46 Pagani, F. & Baralle, F. Pagani, f. & baralle, f. e. genomic variants in exons and introns: identifying the splicing spoilers. nat. rev. genet. 5, 389–396. Nat. reviews. Genet. **5**, 389–96, DOI: 10.1038/nrg1327 (2004).

47 Wang, R., Helbig, I., Edmondson, A. C., Lin, L. & Xing, Y. Splicing defects in rare diseases: transcriptomics and machine learning strategies towards genetic diagnosis. Briefings Bioinforma. 24, bbad284, DOI: 10.1093/bib/bbad284 (2023). https://academic.oup.com/bib/article-pdf/24/5/bbad284/51711356/bbad284.pdf.

48 Varley, J. M. et al. Characterization of germline tp53 splicing mutations and their genetic and functional analysis. Oncogene 20, 2647–2654, DOI: 10.1038/sj.onc.1204369 (2001).

49 Tao, Y., Zhang, Q., Wang, H., Yang, X. & Mu, H. Alternative splicing and related rna binding proteins in human health and disease. Signal transduction targeted therapy 9, 26, DOI: 10.1038/s41392-024-01734-2 (2024).

50 Porter, R., Jaamour, F. & Iwase, S. Neuron-specific alternative splicing of transcriptional machineries: Implications for neurodevelopmental disorders. Mol. Cell. Neurosci. 87, DOI: 10.1016/j.mcn.2017.10.006 (2017).

51 Gandal, M. et al. Transcriptome-wide isoform-level dysregulation in asd, schizophrenia, and bipolar disorder. Science 362, eaat8127, DOI: 10.1126/science.aat8127 (2018).

52 Parras, A. et al. Autism-like phenotype and risk gene mrna deadenylation by cpeb4 mis-splicing. Nature 560, 441–446 (2018).

53 Consortium, T. G. The gtex consortium atlas of genetic regulatory effects across human tissues. Science 369, 1318–1330, DOI: 10.1126/science.aaz1776 (2020).

54 Araujo, P. et al. Before it gets started: Regulating translation at the 5’ utr. Comp. functional genomics 2012, 475731, DOI: 10.1155/2012/475731 (2012).

55 Jackson, R. J., Hellen, C. U. & Pestova, T. V. The mechanism of eukaryotic translation initiation and principles of its regulation. Nat. reviews Mol. cell biology 11, 113–127 (2010).

